# Meta-analysis of GWAS for sea lice load in Atlantic salmon

**DOI:** 10.1101/2022.09.28.509902

**Authors:** P. Cáceres, P. Lopéz, B. Garcia, D. Cichero, J. Ødegård, T. Moen, J.M. Yáñez

## Abstract

Sea lice (*Caligus rogercresseyi*) is an ectoparasite that causes major production losses in the salmon aquaculture industry of the southern hemisphere. Atlantic salmon (*Salmo salar*) is an important salmonid for the aquaculture industry and a species which is highly susceptible to sea lice infestation. Genetic variation for resistance to sea lice, defined as parasite load, has been found in Atlantic salmon. In addition, sea lice load has been shown to be a polygenic trait, controlled by several quantitative trait loci (QTL) which have small to medium effect, making them difficult to map with sufficient statistical power when sample sizes are limited. The use of medium density single nucleotide polymorphisms (SNP) can also adversely affect the success of identifying genetic variants significantly associated to sea lice load. In order to improve the ability to detect QTL significantly associated to sea lice load, we combined genotype imputation from medium- to high SNP-density and performed genome-wide association studies (GWAS) across different populations of Atlantic salmon. The imputation of genotypes of 6,144 fish challenged against sea lice from four year-classes was performed to increase density from 70K SNPs to 600K SNPs. A meta-GWAS was then carried out for three different traits: lice count, lice density and log-lice density. Using this approach, we detected a genomic region highly associated to sea lice load on Atlantic salmon chromosomes (ssa) 3 and 12 pronounced peaks and several other regions surpassing the significance threshold across almost all other chromosomes. We also identified important genes within the QTL regions, many of these genes are involved in tissue reparation, such as Mucin-16-like isoform X2 and Filamentous growth regulator 23-like isoform X1. The QTL region on ssa03 also contained cytoskeletal-modifying and immune response related genes such as Coronin 1A and Claudin. Our results confirm the highly polygenic architecture of sea lice load, but they also show that high experimental power can lead to the identification of candidate genes and thus to increased insight into the biology of sea lice resistance in Altantic salmon.

## 1. Background

Within the animal production sector, aquaculture is the industry that grows most rapidly, and it is expected to have a large impact on the world’s food supply in the next decades [1, 2]. In Chile, the salmon industry was established in the early 1980s. Since then it has grown rapidly and transformed the country into one of the main salmon producing countries the world, harvesting around 435.9 thousand tons of Atlantic salmon (*Salmo salar*) in 2018 [1].

Sea lice is the common name given to a group of ectoparasitic copepods of the family *Caligidae*, which parasitize fish in saltwater [3]. This family accounts for more than 60% of the parasites reported to affect fish in marine environments [4]. The salmon louse, *Lepeophtheirus salmonis*, is only found in the northern hemisphere, predominantly affecting salmonids of the genera *Salmo, Oncorhynchus* and *Salvelinus* [5]. Coho salmon (*Oncorhynchus kisutch*) and pink salmon (*Oncorhynchus gorbuscha*) were described to be more resistant to *L. salmonis* infestation than other salmonid species, such as Atlantic salmon and rainbow trout (*Oncorhynchus mykiss*) [6, 7]. *L. salmonis* was the responsible agent for major salmon disease outbreaks in Canada, Faroe Islands, Ireland, Maine (USA), Norway and Scotland [3, 8]. In the southern hemisphere, *Caligus rogercresseyi* is the main species affecting salmon aquaculture [9]. Infections by *C. rogercresseyi* lead to skin lesions and osmotic imbalance, which in turn increases the salmons susceptibility to secondary bacterial and viral infections [10]. Similarly to *L. salmonis, C. rogercresseyi* seems to affect Atlantic salmon and rainbow trout more than it affects coho salmon, as the latter displays a higher ability to rapidly eliminate the parasites during infestation [11] as well as low sea lice burden under field conditions [12].

A common strategy to combat sea lice is through the use of chemical products, but this strategy is generally limited by the generation of resistance to the drugs in the parasite [13]. One of the alternatives that have been proposed for the control of the parasite is selective breeding for decreased parasite load in the host species [2]. Selecting animals that are more resistant to parasites or have higher ability to eliminate them leads to increases in productivity and animal welfare, and consequently to a lessened environmental impact and reduced costs in the industry [14].

Genome-wide association studies (GWAS) can be used to identify DNA markers associated to traits of importance. This method captures the linkage disequilibrium between markers and causative mutations which tend to be inherited together across generations [15]. GWAS has been applied to provide insights into the genetic architecture of several important traits in Atlantic salmon, including sea lice load [16–19]. All these studies demonstrated a polygenic architecture of lice load, with no genome-wide significant association at any chromosome [16, 19], except for a moderate quantitative trait locus (QTL) located in ssa03 which was reported to surpass the significance threshold in a recent study [17]. Previous studies have most likely failed in detecting significantly associated regions because they used a low to moderate number of animals with phenotypes and genotypes, from 1,498 to 1,119 [18]. 2,600 and 1,056 [17], 2,559 and 2,404 [19] and medium density SNP panels.

An effective strategy to increase the statistical power compared to single-population GWAS and to decrease the rate of false positives is to use meta-analysis of GWAS. In general, this approach may be performed on results from studies of independent populations. For instance, summary statistics from single-population GWAS, including p-values, direction of SNP effects, and sample sizes, can be used to compute an overall Z-score and thus to recalculate GWAS statistics for a meta-population [20]. This approach has been shown to increase the power to detect QTL for polygenic traits in humans [21], and it has also been applied in different livestock species, such as dairy cattle [22], pigs [23] and sheep [24]. A very recent study identified a QTL for the ability to adapt to to high plant protein diet by applying meta-analysis of GWAS large yellow croaker, an important aquaculture species [25].

Genotype imputation can be used to increase the density of genotype information in large populations. This approach infers missing marker information for individuals genotyped with low- to medium-density SNP panels by using a reference population genotyped with a high-density SNP panel. To achieve that, the method uses haplotypes that are shared between both populations [26, 27]. Genotype imputation has two main advantages in genomic applications: reduction of genotyping costs (i.e., genotyping at high density is only necessary for the subset of individuals to be used as reference) and increase of accuracy in detection of genomic regions involved in trait variation [28]. The latter is particularly important for GWAS purposes because the inclusion of ultra-dense SNP information may potentially include causative mutations for the phenotype of interest, which in turn can be used to accelerate the rate of genetic gain if the functional variants are incorporated in the genomic prediction of the genetic merit [29, 30].

The objective of this study was to use meta-analysis of GWAS combined with high-density imputed genotypes to increase the statistical power and accuracy to identify both QTLs and candidate genes associated with sea lice load traits in Atlantic salmon.

## 2. Material and Methods

### 2.1 Origin of animals and sea lice challenges

The data used in this study were obtained from different year-classes (2012, 2013, 2016 and 2017) of the AquaGen breeding population, which were challenged against *C. rogercresseyi*. All challenges were performed in the same way. Briefly, all animals passed by an acclimation period of 13 days on average prior to challenge and were gradually transferred to seawater (32 ppm). After transfer, fish were infested with *C. rogercresseyi* copepodites which had been produced from parasite ovigerous females. The fish were infested with a known number of 35 copepodites per animal. After a number of days ranging from 2 to 18, sea lice load was measured by counting parasites, and weight was also recorded (Table 1). Then, a deinfestation process was applied by gradually reducing the salinity until 5 ppm. After that, salinity levels were increased again to 31 ppm and a new infestation procedure was initiated followed by a second lice counting, as described previously. All year-classes were subjected at least to two different timepoints of lice counting and one year-class (2017) was subjected to four measurement timepoints, in two independent challenges, i.e. two measurements each.

**Table 1.**
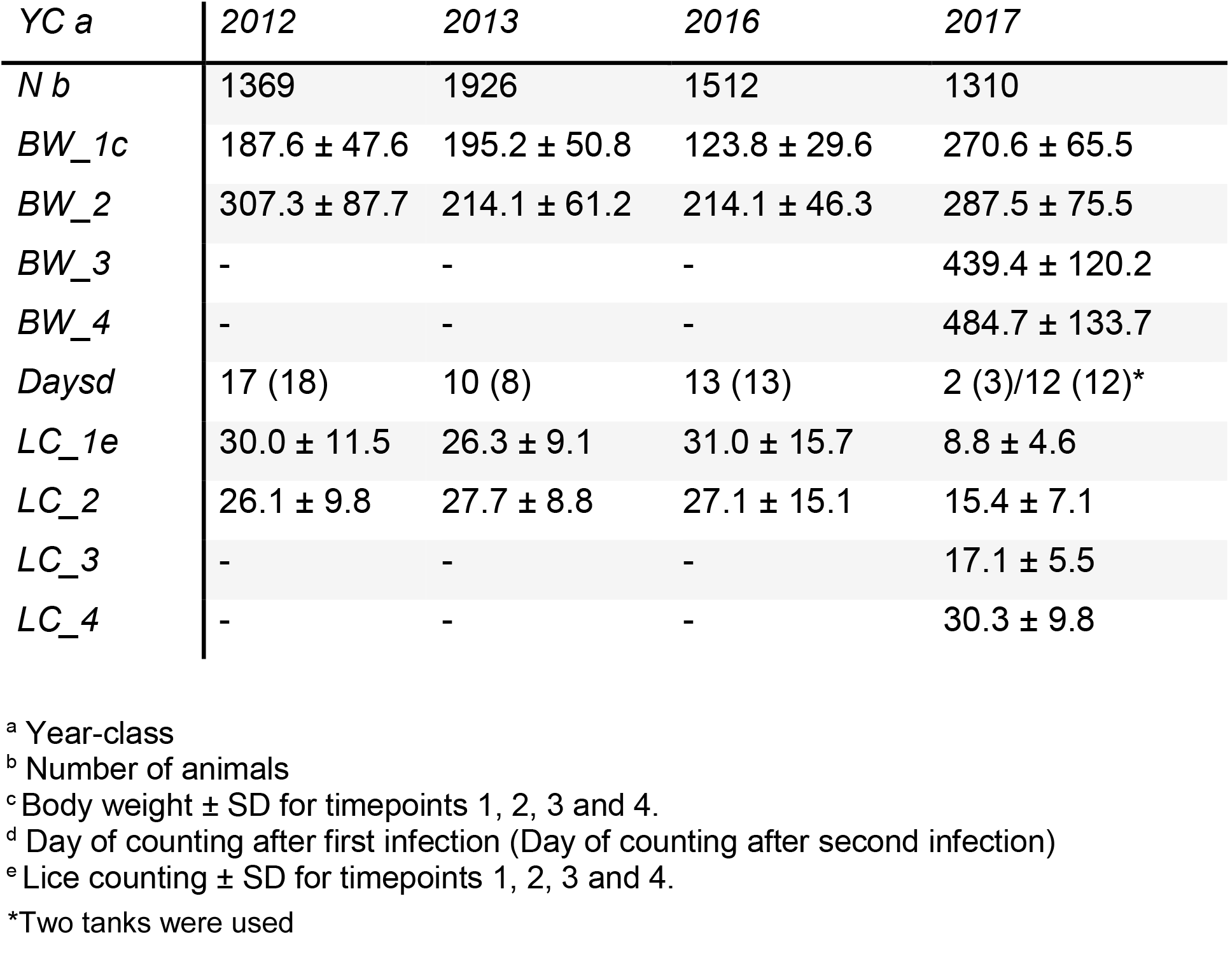
Summary statistics of challenges for sea lice considering number of animals (N), body weight (BW) and lice count (LC) for year-classes at different timepoints.

| YC <sup>a</sup> | 2012 | 2013 | 2016 | 2017 |
| --- | --- | --- | --- | --- |
| N <sup>b</sup> | 1369 | 1926 | 1512 | 1310 |
| BW_1 <sup>c</sup> | 187.6 ± 47.6 | 195.2 ± 50.8 | 123.8 ± 29.6 | 270.6 ± 65.5 |
| BW_2 | 307.3 ± 87.7 | 214.1 ± 61.2 | 214.1 ± 46.3 | 287.5 ± 75.5 |
| BW_3 | - | - | - | 439.4 ± 120.2 |
| BW_4 | - | - | - | 484.7 ± 133.7 |
| Days <sup>d</sup> | 17 (18) | 10 (8) | 13 (13) | 2 (3)/12 (12)* |
| LC_1 <sup>e</sup> | 30.0 ± 11.5 | 26.3 ± 9.1 | 31.0 ± 15.7 | 8.8 ± 4.6 |
| LC_2 | 26.1 ± 9.8 | 27.7 ± 8.8 | 27.1 ± 15.1 | 15.4 ± 7.1 |
| LC_3 | - | - | - | 17.1 ± 5.5 |
| LC_4 | - | - | - | 30.3 ± 9.8 |

<sup>a</sup> Year-class
<sup>b</sup> Number of animals
<sup>c</sup> Body weight ± SD for timepoints 1, 2, 3 and 4.
<sup>d</sup> Day of counting after first infection (Day of counting after second infection)
<sup>e</sup> Lice counting ± SD for timepoints 1, 2, 3 and 4.
\*Two tanks were used

### 2.2 Genotyping

A total of 6,114 animals from all year-classes evaluated in the sea lice challenges were pit-tagged and genotyped using custom ThermoFisher (Affymetrix) SNP arrays developed exclusively for AquaGen’s Atlantic salmon populations. These panels consisted of different versions of medium-density (MD) SNP arrays, containing 50k, 60k and 70k SNP markers, which were used for genomic selection purposes and updated periodically. In order to perform a “step-wise” genotype imputation, we also used genotypic information fromtwo other populations genotyped at 200k and 930k SNP densities (HD_200_ and HD_930_, respectively). The 200k ThermoFisher (Affymetrix) SNP array was used for genotyping 1,480 animals of the 2010 year-class. The 930k ThermoFisher (Affymetrix) SNP array was used for genotyping 1,326 parents of the 2010 year-class and other anteceding animals from previous year-classes [31]. All SNPs used in downstream analyses were positioned based on the Atlantic salmon reference genome (assembly ICSASG_v2).

### 2.3 Quality control and genotype imputation

Initially, a quality control was performed for each genotyped dataset. For all populations genotyped with MD, HD_200_ and HD_930_ SNP arrays, the following filters were applied independently: call-rate for SNPs < 0.9, minor allele frequency (MAF) < 0.01 and Hardy-Weinberg equilibrium (HWE) (Bonferroni corrected p-value < 0.05). **Table 2** shows the number of SNPs before and after quality control for each population included in this study.

**Table 2.**
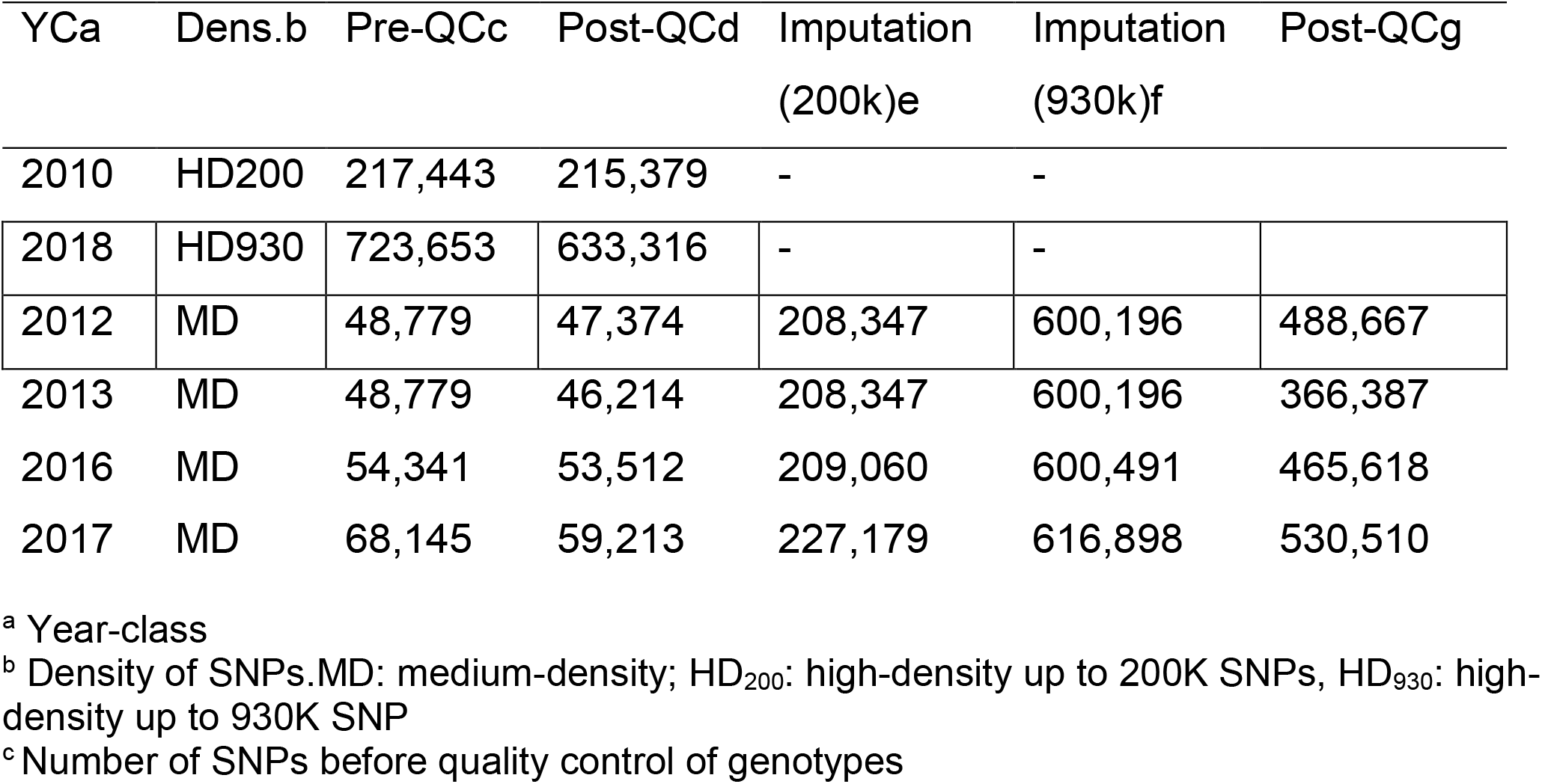
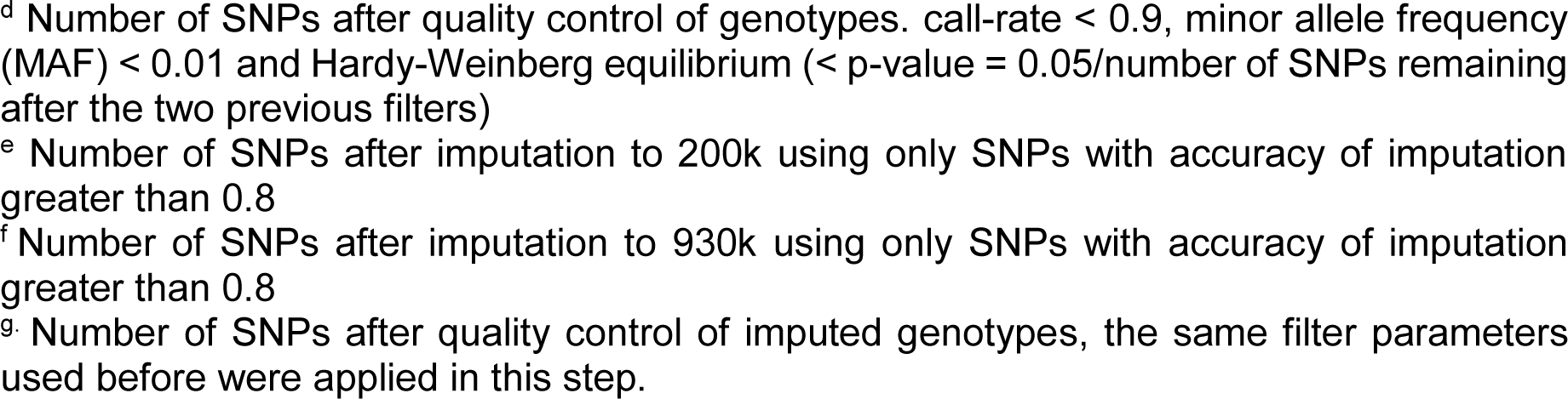
Number of SNPs before and after quality control, and final number of SNPs after imputation to 200k and 930k SNP in each population.

| YCa | Dens.b | Pre-QCc | Post-QCd | Imputation<br>(200k)e | Imputation<br>(930k)f | Post-QCg |
| --- | --- | --- | --- | --- | --- | --- |
| 2010 | HD200 | 217,443 | 215,379 | - | - |  |
| 2018 | HD930 | 723,653 | 633,316 | - | - |  |
| 2012 | MD | 48,779 | 47,374 | 208,347 | 600,196 | 488,667 |
| 2013 | MD | 48,779 | 46,214 | 208,347 | 600,196 | 366,387 |
| 2016 | MD | 54,341 | 53,512 | 209,060 | 600,491 | 465,618 |
| 2017 | MD | 68,145 | 59,213 | 227,179 | 616,898 | 530,510 |

<sup>a</sup> Year-class
<sup>b</sup> Density of SNPs. MD: medium-density; HD<sub>200</sub>: high-density up to 200K SNPs, HD<sub>930</sub>: high-density up to 930K SNP
<sup>c</sup> Number of SNPs before quality control of genotypes
<sup>d</sup> Number of SNPs after quality control of genotypes. call-rate < 0.9, minor allele frequency (MAF) < 0.01 and Hardy-Weinberg equilibrium (< p-value = 0.05/number of SNPs remaining after the two previous filters)
<sup>e</sup> Number of SNPs after imputation to 200k using only SNPs with accuracy of imputation greater than 0.8
<sup>f</sup> Number of SNPs after imputation to 930k using only SNPs with accuracy of imputation greater than 0.8
<sup>g</sup> Number of SNPs after quality control of imputed genotypes, the same filter parameters used before were applied in this step.

A “step-wise” genotype imputation strategy was implemented, imputing first from MD to HD_200_, and then from HD_200_ to HD_930_ as this strategy offers higher accuracy of imputation than direct imputation from medium-density to very high-density [32]. Before genotype imputation, an approximation to imputation accuracy validation was carried out to remove SNPs imputed with low accuracy using a five-folded cross validation scheme for both HD_200_ and HD_930_ dataset. First, 296 animals from the HD_200_ population (20% of total) were randomly assigned as validation population and the other 1,496 animals (80% of total) were assigned as reference population across the five validation groups. The reference animals had 209,579 SNPs and the validation individuals had their genotypes masked keeping only 42,963 SNPs that were in common in the populations genotyped with the MD SNP array. The imputation was performed for each validation group and the accuracy of genotype imputation was estimated using the Pearson’s correlation (r^2^) between imputed and observed genotypes. Finally, only SNPs with mean accuracy of imputation (r^2^) greater than 0.8 were selected to the final set of imputable SNPs. For the HD_930_ validation, the same strategy was implemented using 20% of animals with 633,254 SNPs and 80% of animals 200,394 SNPs as reference and validation populations, respectively. All imputations were performed using the FImpute v3 software using standard parameters [26].

### 2.4 Trait Definitions

To determine the levels of quantitative genetic variation and heritability of sea lice burden, the following trait definitions were used: lice count (LC), lice density (LD) and LogLD. LC was recorded on the skin surface of each individual by manual counting. LD and LogLD were measured accounting for the surface area of the fish, which is determinant for sea lice adherence. Gjerde et al. (2010) defined the lice density (LD) as:

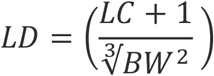

where *BW* is the body weight (g) recorded at the end of the experimental challenge, 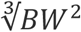 is an approximate measure of the surface area of the fish, and *LC* is the lice count as described previously. In order to normalize the counting data, a transformation to the natural logarithm was applied (LogLD) as:

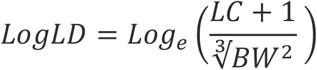

### 2.5 Single-trait GWAS (stGWAS)

The single-trait genome wide association analyses (stGWAS) were performed for each year-class assuming each lice counting at different timepoints, i.e., two records for 2012, 2013 and 2016, and four records for 2017 totalizing 10 records for LC, LD and LogLD each. Prior to stGWAS, a genotype quality control was implemented using Plink software [33], discarding markers with SNP call-rate (< 0.90), MAF (< 0.05) and Hardy-Weinberg Equilibrium (< 0.05 / Number of SNPs). After quality control, the *mlma* option available in the software GCTA v. 1.24 [34] was used to apply the following linear mixed model for each trait:

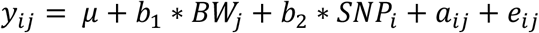

where *y*_*ij*_ is the phenotypic value of the *j*^*-th*^ animal, *μ* is the fixed effect of the overall mean and *b*_1_ and *b*_2_ are regression coefficients and the allele substitution effect for SNP respectively, and *BW*_*j*_ is body weight covariate of *j*^*-th*^ animal and *SNP*_*i*_ is the *i*^*-th*^ SNP genotype of animal *j*, coded as 0, 1 and 2 for genotype A_1_A_1_, A_1_A_2_ and A_2_A_2_, respectively, *a*_*ij*_ is the random polygenic effect of the *j*^*-th*^ animal 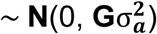, with **G** representing the genomic relationship matrix (GRM) calculated using the imputed genotypes and 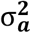 is the genetic variance, and *e*_*ij*_ is the random residual effect 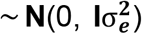, with **I** representing an identity matrix and 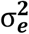 the residual variance. The GRM was calculated based on the relationship between all animals from a genome-wide sample of SNPs obtained by using a common-sense weighting scheme [35]. The GRM restricted maximum likehood (GREML) implemented in GCTA was used to estimate the genetic and residual variances. Heritability (*h*^*2*^) was calculated as 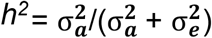. For each SNP, the variance explained was also calculated using the following equation [36]:

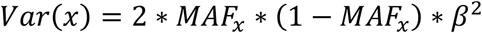

where *Var*(*x*) is the variance explained by the SNP *x, MAF*_*x*_ is the minor allele frequency of SNP*x* and *β* is the SNP*x* effect.

### 2.6 Meta-analysis of stGWAS results (metaGWAS)

For each trait (LC, LD and LogLD), a meta-analysis of GWAS results using all year-classes was performed using the METAL software [20]. The p-value, direction of effect, and sample size were utilized to implement a sample size weighted analysis, with additional genomic control collection performed based on the difference between the median test statistic and that expected by chance computing an overall Z-score:

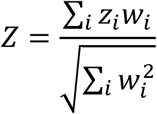

where *w*_*i*_ is the square root of the sample size of population *i*, and *z*_*i*_ is the Z-score for population *i* calculated as 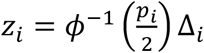, where *ϕ* is the cumulative distribution function, and *p*_*i*_ and Δ_*i*_ are the p-value and direction of effect for population *i*, respectively. Manhattan plots were produced for each trait in order to visualize the results from the meta-analyses.

### 2.7 Identification of QTL and candidate genes

Lead SNPs were selected based on a genomic threshold with Bonferroni correction (0.05 / number of SNPs). All variants with p-value crossing the threshold, were considered a lead SNP, and used to search for candidate genes based on proximity to the variant. The gene search was performed using BLAST (Basic Local Alignment Search Tool) against the *Salmo salar* reference genome (ICSASG_v2), which is publicly available at NCBI (GenBank assembly accession GCA_000233375.4). Genes located within 100 kb upstream and downstream of the leading SNPs were considered putative candidate genes associated with the trait.

## 3. Results

### 3.1 Descriptive statistics

Summary statistics from each sea lice challenge test, including number of animals, mean body weight ± sd, day of counting post infection and LC ± sd are shown in Table 1. The differences in days of counting among the challenges can be explained by several attempts to understand the kinetic of infestation, mainly focused on providing a better understanding on the lice attachment process. However, all LC measures were very similar across different year-classes, except for the 2017 in which an early time-point was included as the first the counting day (two and three days post infestation).

### 3.2 Genotype imputation

A total of 167,751 and 393,157 additional SNPs with r^2^ greater than 0.8 were obtained after imputation from MD to HD_200_ and HD_930_, respectively. Adding up the SNPs originally genotyped and the imputed SNPs, it was possible to achieve approximately 600k SNPs as the final density (Table 2). For all imputed HD_200_ and HD_930_ genotypes, quality filters were also applied for each population separately. For SNP call-rate (< 0.9), only 2017 year-class had markers removed (3,119 SNPs). MAF < 0.05 was the filter that discarded most SNPs as 111,972 (18.4%), 192,623 (31.7%), 136,896 (22.5%) and 131,951 (21.2%) were removed for 2012, 2013,2016 and 2017 year-classes, respectively. The HWE filter discarded a lower number of markers, 7,453 (1.5%), 49,040(11.8%), 6,170 (1.3%) and 5,165 (1.0%) for 2012, 2013, 2016 and 2017 year-classes, respectively. The final number of SNPs after quality control before running stGWAS was 488,677, 366,387, 465,618 and 530,510 for 2012, 2013, 2016 and 2017 year-classes, respectively.

### 3.3 Genetic parameters

Heritability estimates calculated using the SNP-based GRM constructed with about ∼500.000 markers for each population ranged from 0.06 ± 0.02 to 0.23 ± 0.04 for LC, 0.02 ± 0.01 to 0.29 ± 0.04 for LD and 0.10 ± 0.03 to 0.36 ± 0.03 for LogLD as show in Table 3.

**Table 3.** Genetic parameters of every year class for all traits evaluated in the meta-analysis.

| YC<sup>a</sup> | Trait<sup>b</sup> | V(e)<sup>c</sup> | V(g)<sup>d</sup> | h<sup>2e</sup> |
| --- | --- | --- | --- | --- |
| 2012 | LC_18 | 118.70 ± 5.38 | 7.73 ± 3.49 | 0.06 ± 0.02 |
|  | LC_18 | 79.78 ± 3.89 | 17.41 ± 4.02 | 0.17 ± 0.03 |
|  | LD_18 | 0.12 ± 0.005 | 0.01 ± 0.004 | 0.11 ± 0.03 |
|  | LD_18 | 0.05 ± 0.003 | 0.02 ± 0.004 | 0.29 ± 0.04 |
|  | LogLD_18 | 0.13 ± 0.006 | 0.01 ± 0.005 | 0.10 ± 0.03 |
|  | LogLD_18 | 0.14 ± 0.007 | 0.04 ± 0.009 | 0.24 ± 0.04 |
| 2013 | LC_10 | 56.42 ± 2.30 | 12.91 ± 2.47 | 0.18 ± 0.03 |
|  | LC_8 | 62.62 ± 2.35 | 6.53 ± 1.78 | 0.09 ± 0.02 |
|  | LD_10 | 0.06 ± 0.002 | 0.01 ± 0.002 | 0.15 ± 0.03 |
|  | LD_8 | 0.05 ± 0.002 | 0.006e<sup>-</sup> ± 0.001 | 0.10 ± 0.02 |
|  | LogLD_10 | 0.20 ± 0.009 | 0.084 ± 0.012 | 0.28 ± 0.03 |
|  | LogLD_8 | 0.20 ± 0.009 | 0.11 ± 0.01 | 0.36 ± 0.03 |
| 2016 | LC_13 | 187.38 ± 8.61 | 45.09 ± 9.014 | 0.19 ± 0.03 |
|  | LC_13 | 177.26 ± 8.07 | 38.50 ± 8.11 | 0.17 ± 0.03 |
|  | LD_13 | 0.31 ± 0.01 | 0.07 ± 0.01 | 0.19 ± 0.03 |
|  | LD_13 | 0.14 ± 0.006 | 0.03 ± 0.006 | 0.18 ± 0.03 |
|  | LogLD_13 | 0.22 ± 0.01 | 0.095 ± 0.015 | 0.29 ± 0.03 |
|  | LogLD_13 | 0.25±0.011 | 0.06 ± 0.012 | 0.19 ± 0.03 |
| 2017 | LC_2 | 28.59 ± 1.49 | 6.90 ± 1.56 | 0.19 ± 0.04 |
|  | LC_12 | 59.45 ± 3.15 | 18.29 ± 3.56 | 0.23 ± 0.04 |
|  | LC_3 | 45.80 ± 2.31 | 9.16 ± 2.22 | 0.16 ± 0.03 |
|  | LC_12 | 103.51 ± 5.48 | 30.58 ± 6.15 | 0.22 ± 0.04 |
|  | LD_2 | 0.01 ± 0.001 | 0.004 ± 9e-4 | 0.19 ± 0.04 |
|  | LD_12 | 0.03 ± 0.001 | 0.01 ± 0.002 | 0.22 ± 0.04 |
|  | LD_3 | 0.014 ± 7e-4 | 0.002 ± 7e-4 | 0.16 ± 0.03 |
|  | LD_12 | 0.03 ± 0.001 | 0.008 ± 0.001 | 0.22 ± 0.04 |
|  | LogLD_2 | 0.32 ± 0.017 | 0.08 ± 0.018 | 0.21 ± 0.04 |
|  | LogLD_12 | 0.24 ± 0.012 | 0.06 ± 0.013 | 0.21 ± 0.03 |
|  | LogLD_3 | 0.12 ± 0.006 | 0.02 ± 0.005 | 0.15 ± 0.03 |
|  | LogLD_12 | 0.11 ± 0.006 | 0.032 ± 0.006 | 0.21 ± 0.04 |

<sup>a</sup> Year-class
<sup>b</sup> Trait. LC: lice count; LD: lice density; LogLD: Log of lice density. Numbers after the underscore mean the correlative counting timepoint during the challenge tets (from 1 to 4).
<sup>c</sup> Residual variance ± SD
<sup>d</sup> Genetic variance ± SD
<sup>e</sup> Heritability ± SD

### 3.4 stGWAS and metaGWAS

For all stGWAS in each population, there were only few SNPs that surpassed the genome-wide significance threshold, no major quantitative trait loci were identified on each particular population when running stGWAS. For LC, the lowest p-value was observed at 18 day post infestation of the 2012 year-class on ssa17 (p-value=1.25e-^07^). For LD and LogLD, the lowest p-values were also observed at 18 day post-infestation of the 2012 year-class, but in this case on ssa26 (p-value=7.52e^-09^ and 1.23e^-08^, respectively). The Manhattan plots of stGWAS for every year-class and trait are shown in Supplementary Figure 1. Few SNPs explaining between 1 to 4 percent of the fraction of additive genetic variance (Supplementary Table 1).

For the metaGWAS, a substantial increase in the significance level for SNP-trait association was observed on all chromosomes (Figure 1). A pronounced significant peak was identified for all traits on ssa03, with p-values of 1.85e^-22^, 9.75e^-24^ and 2.45e^-31^ for LC, LD and LogLD, respectively. Smaller consistent peaks were also observed on ssa12 (p-values of 1.10e^-19^, 8.79e^-20^ and 5.73e^-19^ for LC, LD and LogLD, respectively), ssa18 (p-values of 5.71e-^15^, 1.11^e-12^ and 4.45^e-14^ for LC, LD and LogLD, respectively) and ssa22 (p-values of 7.928e^-16^, 2.356^e-15^ and 1.060^e-12^ for LC, LD and LogLD, respectively). In addition, it is important to mention that significant markers were widely distributed across almost all the chromosomes for all traits. These results highlight the polygenic architecture of sea lice load in Atlantic salmon, but also show that potential QTLs with minor to moderate effects may be also affecting these traits.

**Figure 1.**
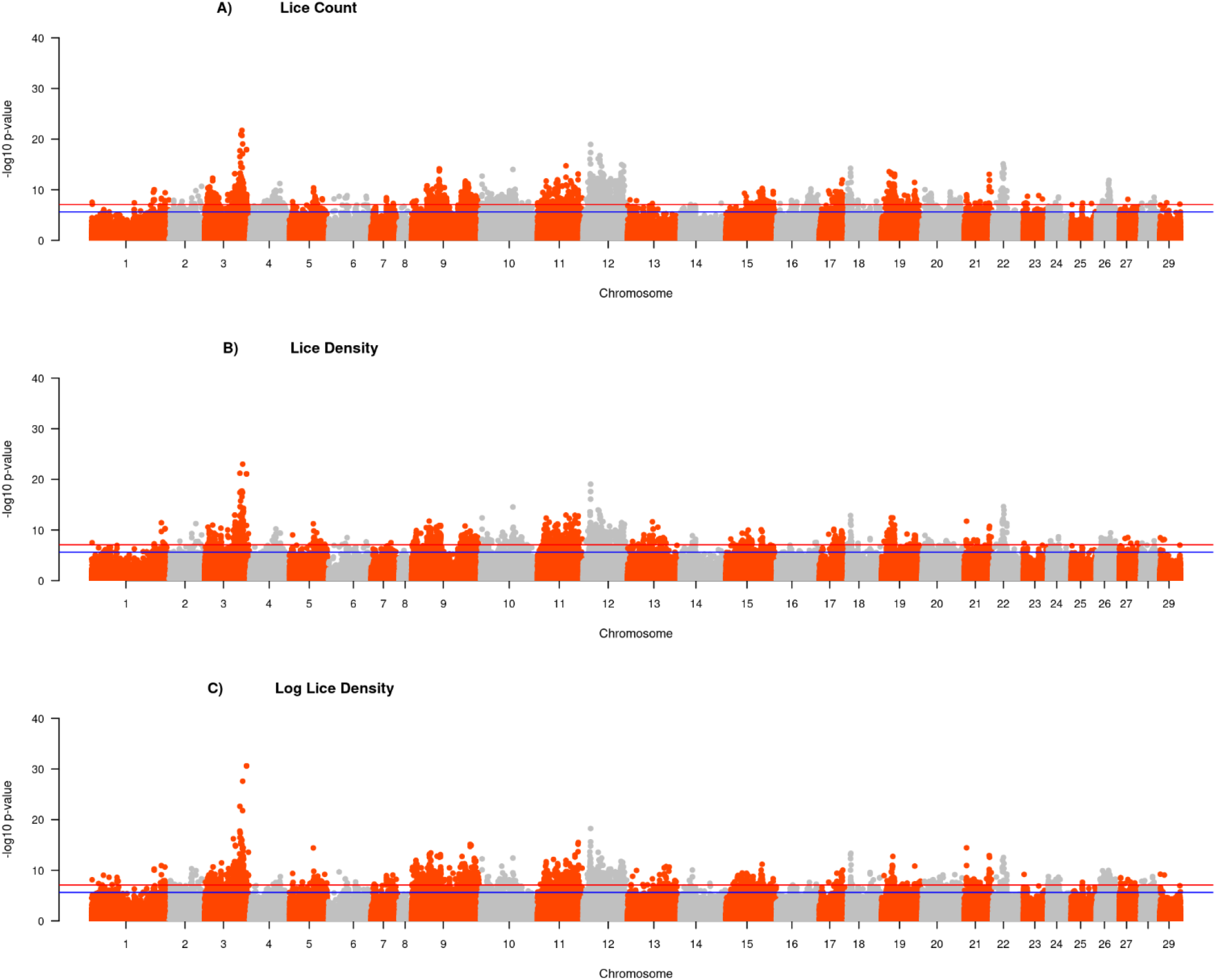
Manhattan plots of meta-analysis of GWAS for lice count (A), lice density and Logarithm of lice density (C) traits. The blue and red lines indicate the chromosome- and genome-wide significance thresholds, respectively.

### 3.5 Candidate genes

A summary of the main top markers found in the QTL regions are in Table 4, and the full list of genes located within 100 kb upstream and downstream of the lead SNP is available in Supplementary Table 2. Some lead SNPs for LC are close to important candidate genes, including Mucin-16-like isoform X2 and Filamentous growth regulator 23-like isoform X1 on ssa03, and Fibroblast growth factor receptor-like 1 on ssa12. Some of these genes were already associated with sea lice load variation in previous studies performed in an independent population in Atlantic salmon [17].

**Table 4.**
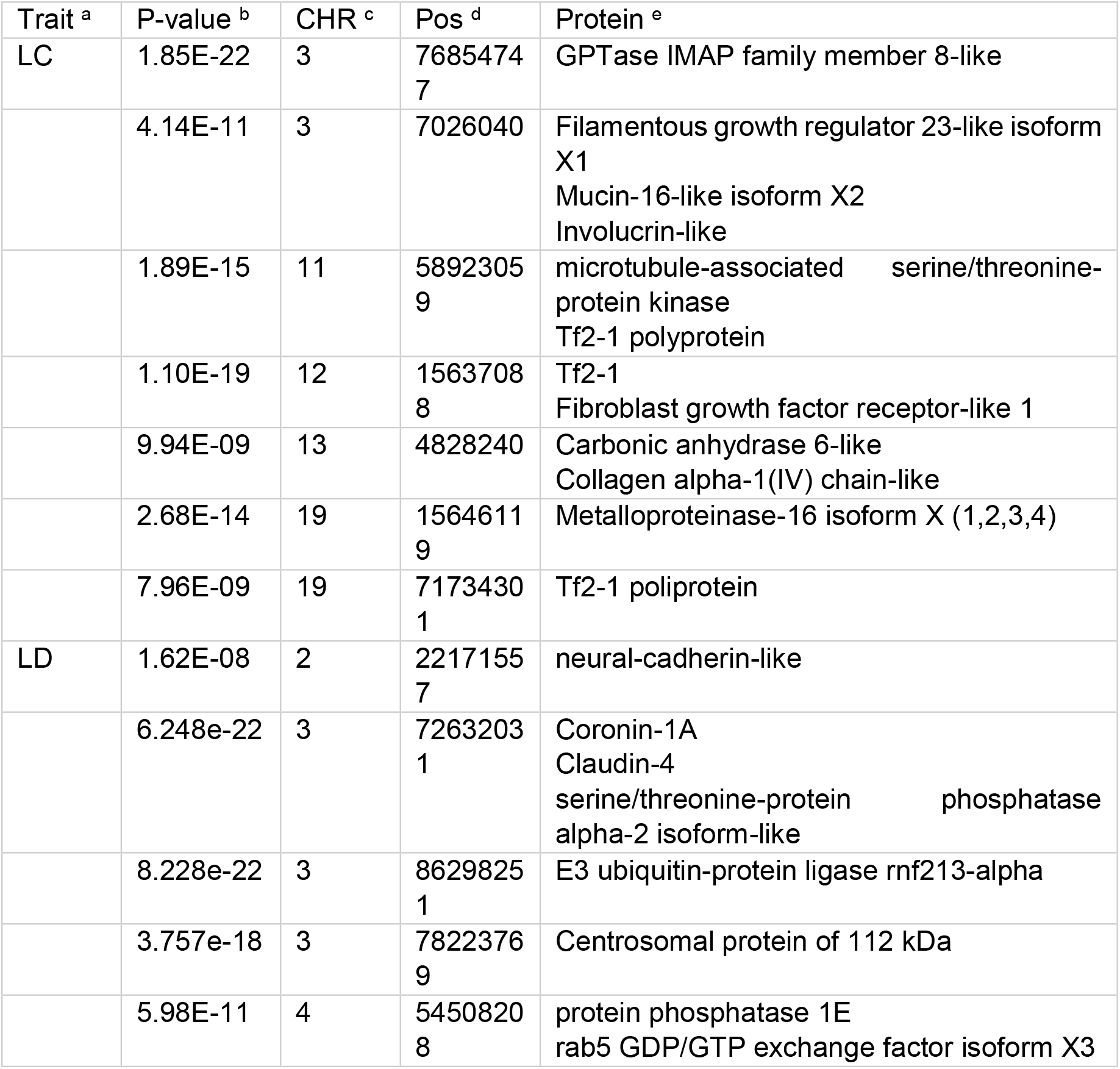
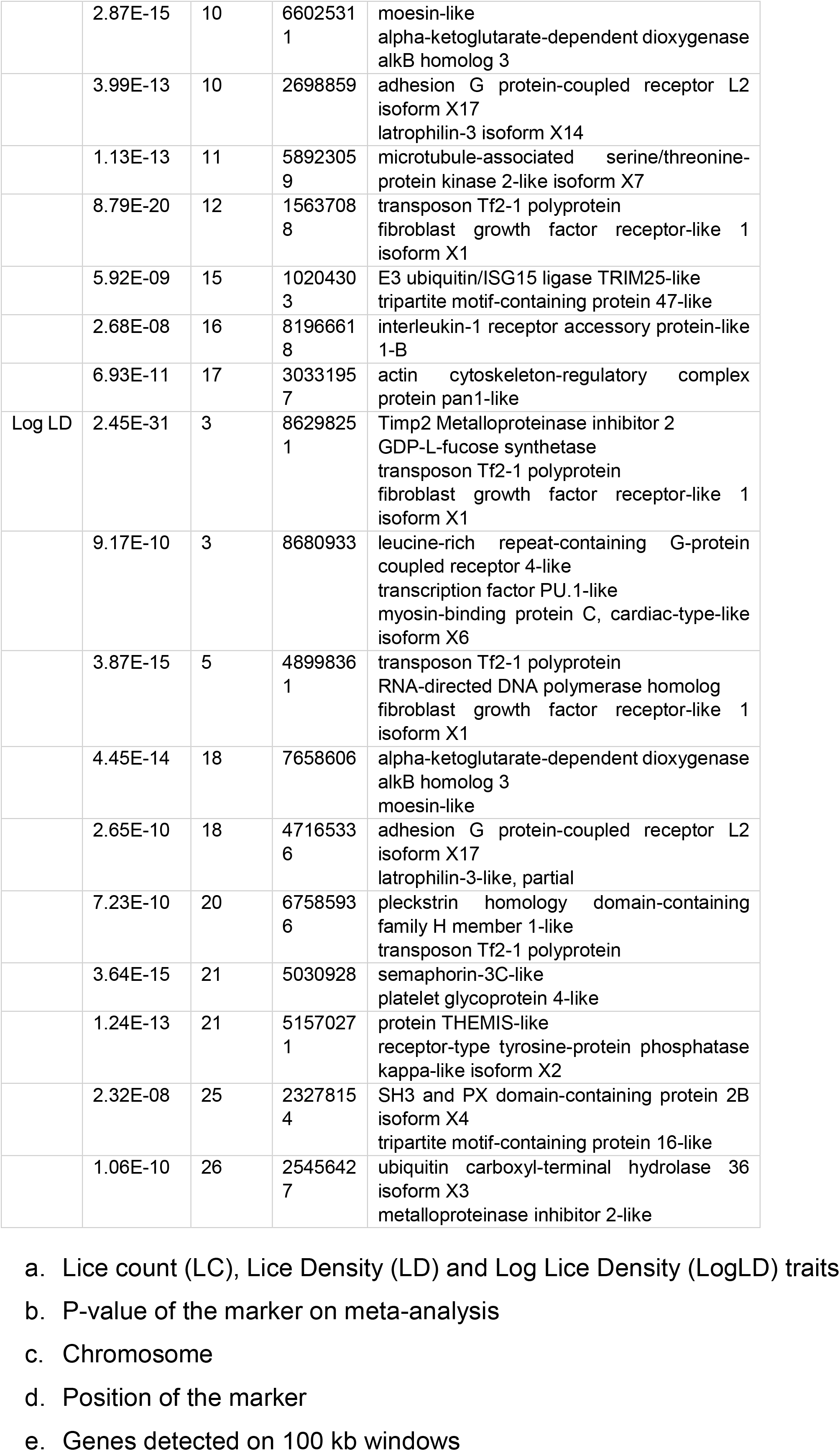
Summary of top markers associated with sea lice traits on meta-analysis.

For LD, a lead SNP was found near to a Coronin 1A and Claudin 4 genes between 72,636,574 and 72,647,185 bp position on ssa03. Both genes are related to the immune system and cytoskeleton modification. The same SNP at ssa03 overlapped the serine/threonine-protein phosphatase alpha-2 isoform-like gene, which was also associated with sea lice resistance on GWAS and functional studies in Atlantic salmon [19, 37]. Additionally in the same chromosome, other 2 SNPs intercepted an exonic region of the centrosomal protein of 112 kDa at 78,079,260 bpp, which is related to microtubules formation on mammalian cells [38]. Other lead SNPs were found on ssa10 at 2,632,647 bp, overlapped the gene of GTP binding protein 6, and on ssa12 at 15,513,054 bp near to an exonic region of the alpha-2-macroglobulin-like gene, which is a proteinase regulator [39].

For LogLD, a lead SNP overlapped the gene of metalloproteinase inhibitor 2 (*timp2*) at 86,314,119 bp on ssa03 and is nearby to GDP-L-fucose synthetase gene. The metalloproteinases inhibitor may participate in a wide range of processes, such as activation and release of cytokines, inhibition of growth factors from extracellular matrix and favoring of cells migration to the wound area [40]. Another lead SNP on the same chromosome is close to somatostatin receptor type 5-like at 72,684,148 bp. On ssa09, a lead SNP is nearby E3 ubiquitin-protein ligase (*trim*8), which was identified as a candidate gene for sea lice resistance in a previous study [19]. A lead SNP on ssa12 intercepted an exonic region of ral guanine nucleotide dissociation stimulator-like 1 at 15,628,486 bp and, at the same chromosome, a SNP is near to the filaggrin-2-like gene that participates in cell to cell adhesion and structure of epidermis [41].

## 4. Discussion

We found low to moderate (0.02 ± 0.01 to 0.36 ± 0.03) heritabilities for LC, LD and LogLD traits considering all data sets. These results agree with previous studies in which sea lice resistance is defined as parasite burden. Previous studies have estimated heritabilities using genomic information with SNP panel densities from 60K to 200K [10, 17, 19, 42–48]. The use of imputed information increased heritability values for body weight traits as was observed in Nile tilapia [49, 50] using the same population from medium to ultra high density and for sea lice trait [51], using low density to high density imputed genotypes.

Previous studies identified loci and candidate genes associated with sea lice resistance traits in Atlantic salmon [17, 19, 52], but the medium-density SNP panels and limited sample size used in these studies may have hindered the identification of genes truly associated to these traits. These GWAS were performed with moderate sample size using a maximum of ∼2,600 animals, and using only medium density SNP panels[10, 16, 18]..

In order to increase stadistical power and accuracy for detecting association between SNPs and the traits of interest, we imputed genotypes to high-density and then performed meta-analysis of GWAS. The imputation was performed from medium-density SNP arrays (50, 60 and 70K) to high-density at two different levels, ∼200K and ∼500K, using markers previously selected based on expected accuracy of imputation higher than 0.80. The genotype imputation may help to increase GWAS resolution allowing fine-mapping of the sea lice load traits. However, as shown for stGWAS for all year-classes (Supplementary Figure 1), few SNPs surpassed the genome-wide significance threshold, similar to previous GWAS results for the same traits performed in independent populations, in which no evidence of major effect QTLs for sea lice load traits was found [19].

The metaGWAS effectively increased significance of genomic regions identified in our study. This approach was applied to leverage multiple summary statistics from stGWAS performed on the same traits using different populations (year-classes) and increasing the sample size. In addition to increasing sample size, the metaGWAS strategy helps to reduce the number of false positives because most of effect sizes usually detected in GWAS are small and inconsistent across different studies or populations [53].

The difference in the number of genome-wide statistically significant markers associated with sea lice load traits between stGWAS and metaGWAS was remarkable. Overall, the increase in significant SNPs was consistent across all traits, for LC, LD and Log LD we found 2,340, 1,872 and 2,204 SNPs who cross the genomic significant threshold, respectively. However, there were also significant SNPs in the stGWAS analyses that were not confirmed in the metaGWAS. If the SNP association is not confirmed in the metaGWAS, we may assume that the prior association identified by GWAS is only relevant to the year-class in which is detected or might be a false positive.

We found approximately 2,000 genome-wide significant markers dispersed in almost all chromosomes, and these markers may be tagging important genetic variants controlling sea lice load in Atlantic salmon. This result was expected given that sea lice load has been previously described as a polygenic trait in this species [43, 52, 54]. A pronounced peak comprised by highly significant markers on ssa03 was found for all traits analyzed, suggesting that part of the genetic variation in sea lice load is due to a low to moderate effect QTL on this chromosome. This region on ssa03 is consistent with previous results from GWAS performed for similar traits in different populations [19, 37].

The lead SNP in ssa03 is nearby the metalloproteinase inhibitor 2 (*timp*2). Several metalloproteinases were associated with functional response to sea lice infestation [40, 55, 56], participating in important innate response processes including inflammation and tissue remodeling. Within the same region, another lead SNP is near to cytohesin-1 (*cyh*1) gene which is an abundant protein of immune cells, participating on cell binding and adhesion [57]. A related protein potentially associated to sea lice response, the serine/threonine-protein phosphatase alpha-2 isoform-like, was also detected on ssa03, and described to be playing an important role in cell growth and signaling [58]. A comparative transcriptomic analysis between healthy skin and skin where sea lice was attached in Atlantic salmon showed significant differences on the expression ratio of this protein [40, 54]. Genes related to this protein were also identified in GWAS for sea lice load traits in an experimentally challenged rainbow trout population [19].

At the same chromosome, we identified novel candidate genes, such as Claudin and Coronin between 72,636,574 and 72,647,185 bp. The SNPs close to these genes are strongly associated with sea lice load. Claudins are cell-to-cell adhesion molecules located at the tight junctions between cells in epithelial cell sheets, creating a physical barrier against the external environment [59]. The tight junctions between epithelial cells act as a selective permeable barrier that regulates the movement of solutes between fluid compartments, thus, they are important determinants of ion selectivity and general permeability of the epithelia [60]. The Coronin is a conserved actin binding protein that promotes cellular processes that rely on rapid remodeling of the actin cytoskeleton, including endocytosis and cell motility [61].

Significant SNPs were also found at different chromosomes close to widely characterized genes involved in tissue repair, such as the Fibroblast growth factor receptor-like 1 on ssa12 [62], Collagen alpha-1 (IV) chain-like on ssa13 and Collagen alpha-6 (IV) chain-like [63] on ssa18. These tissue repair genes were described in genomics and functional studies focused on host response against sea lice infestation and have great importance in tissue reaction to the parasite in coho salmon, a species considered highly resistant to sea lice infestation [64]. Collagen and fibroblast growth factor genes could be related to the response of fish to skin wounds, as described by [65]. These molecules are involved in processes of proliferation of filamentous cells and sealing the cutaneous wound, followed by leukocyte infiltration and epidermal thickening and hyperplasia[11]. The skin wound maturation occurs through increased epidermal layers and thickness and migration of cells rather than proliferation of cells [65]. After the inflammation, a concerted action of different cell types occurs, all coordinated by a complex network of growth factors and other regulators. Enzymes, such as metalloproteinases [66], are able to destroy components of the extracellular matrix that are involved in both inflammation and tissue repair.

Although our study provides valuable insights about genomic regions associated to sea lice resistance in Atlantic salmon, it is still unclear how the whole mechanism of defense works for this species.. Coho salmon, a species considered highly resistant to sea lice, shows quick inflammatory response and epithelial hyperplasia generating an efficient mechanism that may reduce up to 90% of lice loads in two weeks [11]. The inflammatory response seems to be mild or incomplete in Atlantic salmon with low neutrophils mobilization at the site of infection [67]. For this reason, understanding how the infection is fought in resistant species (e.g., coho salmon) may help to unravel the high susceptibility of other species (e.g., Atlantic salmon and rainbow trout).

It is also important to mention that all challenges were performed in controlled conditions. There is a possibility that under field conditions, such as with sea cages, the response mechanisms may act or interact distinctly to controlled conditions given by experimental challenges. Another hypothesis is that, under controlled conditions, host factors (e.g., susceptibility, attractiveness, selectivity) [68] are being evaluated instead of the true ability of hosts to eliminate the parasites. For this reason, more studies regarding the kinetics of sea lice in Atlantic salmon under sea conditions are needed for distinguishing the different host and environmental factors affecting sea lice load, from the actual variation of individual response against the infection.

Still, the genes found in the present study, which are putatively associated to the variation of response to sea lice infestation in Atlantic salmon, confirms that inflammatory, cell mobilization and tissue repair processes are biological functions that could be determinant on individual variation of sea lice burden in this species [56, 69].

## Conclusion

The meta-analysis using genotypes imputed to high-density helped to detect SNPs significantly associated to sea lice load in Atlantic salmon. Our results confirm that sea lice load related traits are polygenic in nature. Nevertheless, we found several putative candidate genes which may be contributing to the genetic variance of sea lice load in this species. Tissue repair, cytoskeletal modification and immune response ay be playing an important role in the individual variation of sea lice burden in experimentally challenged Atlantic salmon populations.

## Acknowledgments

JMY would like to thank funding from FONDECYT Regular (No. 1211761) and Millennium Science Initiative NCN2021_056: Millennium Nucleus of Austral Invasive Salmonids, funded by Chile’s Government Programme, Ministerio de Economia, Fomento y Turismo, PC is supported by doctoral funding (No. 21211569) of Agencia Nacional de Investigación y Desarrollo (ANID).

## Author contributions

PC. assessed the analyses and wrote the initial version of the manuscript. PL, DC and TM. managed samples, performed DNA extraction and performed the quality control of genotypes. BG and JØ. contributed with writing, revision and discussion of the results, JMY conceived the study, contributed to results interpretation with a discussion. All authors approved the manuscript.

