## Supplementary Table 1 for "Meta-analysis of GWAS for sea lice load in Atlantic salmon"

| Chr | SNP | Bpp | Year_class | Trait | b_effect |
|---|---|---|---|---|---|
| 26 | ctg7180001524632_9188_SGT | 15113524 | SS12 | logld2 | 0.147197 |
| 26 | ctg7180001855138_1619_SCT | 14952628 | SS12 | logld2 | 0.119585 |
| 26 | ctg7180001291339_103_SAG | 14954394 | SS12 | logld2 | 0.119585 |
| 26 | ctg7180001291339_114_SAT | 14954405 | SS12 | logld2 | 0.119585 |
| 26 | ctg7180001803978_10943_SAC | 14763442 | SS12 | logld2 | 0.135872 |
| 13 | ctg7180001902113_3711_SGT | 89439192 | SS12 | logld2 | 0.109433 |
| 17 | ctg7180001868430_1021_SAG | 51584064 | SS12 | logld2 | 0.12515 |
| 9 | ctg7180001926755_8001_SCT | 115780139 | SS12 | logld2 | 0.082704 |
| 20 | ctg7180001937452_12443_SAC | 8125754 | SS12 | logld2 | 0.096949 |
| 26 | scf1519059855654T | 15171760 | SS12 | logld2 | 0.086425 |
| 13 | ctg7180001903381_1649_SCT | 91262398 | SS12 | logld1 | 0.089373 |
| 13 | ctg7180001903381_1791_SAG | 91262540 | SS12 | logld1 | 0.08736 |
| 13 | ctg7180001903381_1959_SCT | 91262708 | SS12 | logld1 | 0.08736 |
| 13 | ctg7180001903381_6930_SCT | 91267679 | SS12 | logld1 | 0.08736 |
| 13 | ctg7180001898975_1435_SGT | 95648119 | SS12 | logld1 | 0.083905 |
| 13 | ctg7180001730411_3854_SGT | 95653384 | SS12 | logld1 | 0.083905 |
| 13 | ctg7180001908796_14602_SAC | 91338281 | SS12 | logld1 | 0.081561 |
| 13 | ctg7180001822551_4707_SAG | 91289206 | SS12 | logld1 | 0.080934 |
| 2 | ctg7180001725809_1994_SAG | 23294888 | SS12 | logld1 | 0.074267 |
| 24 | ctg7180001376884_107_SGT | 45414276 | SS12 | logld1 | -0.11346 |
| 26 | ctg7180001524632_9188_SGT | 15113524 | SS12 | ld2 | 0.097324 |
| 17 | ctg7180001868430_1021_SAG | 51584064 | SS12 | ld2 | 0.093163 |
| 26 | scf1519059855654T | 15171760 | SS12 | ld2 | 0.064288 |
| 26 | ctg7180001855138_1619_SCT | 14952628 | SS12 | ld2 | 0.080925 |
| 26 | ctg7180001291339_103_SAG | 14954394 | SS12 | ld2 | 0.080925 |
| 26 | ctg7180001291339_114_SAT | 14954405 | SS12 | ld2 | 0.080925 |
| 13 | ctg7180001902113_3711_SGT | 89439192 | SS12 | ld2 | 0.077196 |
| 20 | ctg7180001916753_8656_SGT | 13319314 | SS12 | ld2 | 0.125592 |
| 26 | ctg7180001793181_2093_SAG | 16108919 | SS12 | ld2 | 0.069967 |
| 17 | ctg7180001795677_6478_SAC | 51548695 | SS12 | ld2 | 0.088307 |
| 17 | ctg7180001795677_6532_SAG | 51548749 | SS12 | ld2 | 0.088307 |
| 13 | ctg7180001903381_1649_SCT | 91262398 | SS12 | ld1 | 0.085869 |
| 13 | ctg7180001903381_1791_SAG | 91262540 | SS12 | ld1 | 0.083782 |
| 13 | ctg7180001903381_1959_SCT | 91262708 | SS12 | ld1 | 0.083782 |
| 13 | ctg7180001903381_6930_SCT | 91267679 | SS12 | ld1 | 0.083782 |
| 13 | ctg7180001898975_1435_SGT | 95648119 | SS12 | ld1 | 0.079563 |
| 13 | ctg7180001730411_3854_SGT | 95653384 | SS12 | ld1 | 0.079563 |
| 13 | ctg7180001908796_14602_SAC | 91338281 | SS12 | ld1 | 0.07781 |
| 13 | ctg7180001822551_4707_SAG | 91289206 | SS12 | ld1 | 0.077299 |
| 13 | ctg7180001918104_3421_SAG | 87342639 | SS12 | ld1 | -0.07566 |
| 2 | ctg7180001912875_1559_SCG | 7101162 | SS12 | ld1 | 0.074997 |
| 17 | ctg7180001868430_1021_SAG | 51584064 | SS12 | lc2 | 3.3605 |
| 17 | ctg7180001934232_8785_SGT | 43627523 | SS12 | lc2 | 3.82606 |
| 17 | ctg7180001795677_6478_SAC | 51548695 | SS12 | lc2 | 3.19939 |
| 17 | ctg7180001795677_6532_SAG | 51548749 | SS12 | lc2 | 3.19939 |
| 17 | ctg7180001395898_1445_SCT | 52254328 | SS12 | lc2 | 3.09982 |
| 17 | ctg7180001395898_1994_SAG | 52254877 | SS12 | lc2 | 3.09982 |
| 17 | ctg7180001925542_3041_SGT | 52284839 | SS12 | lc2 | 2.85023 |
| 17 | ctg7180001807695_2808_SAG | 52309721 | SS12 | lc2 | 2.85023 |

| | | | | | |
|---|---|---|---|---|---|
| 17 | ctg7180001809166_3615_SAG | 50940729 | SS12 | lc2 | 2.87143 |
| 17 | ctg7180001818203_683_SAC | 52368362 | SS12 | lc2 | 2.98655 |
| 13 | ctg7180001903381_1649_SCT | 91262398 | SS12 | lc1 | 2.40873 |
| 13 | ctg7180001903381_1791_SAG | 91262540 | SS12 | lc1 | 2.33684 |
| 13 | ctg7180001903381_1959_SCT | 91262708 | SS12 | lc1 | 2.33684 |
| 13 | ctg7180001903381_6930_SCT | 91267679 | SS12 | lc1 | 2.33684 |
| 13 | ctg7180001898975_1435_SGT | 95648119 | SS12 | lc1 | 2.22194 |
| 13 | ctg7180001730411_3854_SGT | 95653384 | SS12 | lc1 | 2.22194 |
| 2 | ctg7180001912875_1559_SCG | 7101162 | SS12 | lc1 | 2.20053 |
| 2 | ctg7180001544546_2120_SAG | 7007689 | SS12 | lc1 | 2.25161 |
| 2 | ctg7180001912875_6284_SAC | 7096435 | SS12 | lc1 | 2.25161 |
| 13 | ctg7180001805890_9840_SAG | 65700702 | SS12 | lc1 | 2.29463 |
| 13 | ctg7180001805890_9129_SAG | 65701413 | SS12 | lc1 | 2.29463 |
| 3 | ctg7180001825788_12570_SAC | 70193555 | SS13 | logld2 | -0.07721 |
| 26 | ctg7180001809454_9640_SGT | 11133973 | SS13 | logld2 | 0.064257 |
| 26 | ctg7180001809454_6285_SCT | 11137328 | SS13 | logld2 | 0.064257 |
| 26 | ctg7180001809454_6259_SAC | 11137354 | SS13 | logld2 | 0.064257 |
| 26 | ctg7180001809454_6191_SAG | 11137422 | SS13 | logld2 | 0.064257 |
| 26 | ctg7180001809454_6104_SAG | 11137509 | SS13 | logld2 | 0.064257 |
| 26 | ctg7180001809454_3127_SAC | 11140486 | SS13 | logld2 | 0.064257 |
| 26 | ctg7180001809454_2266_SAC | 11141347 | SS13 | logld2 | 0.064257 |
| 9 | ctg7180001361659_1650_SAG | 6064750 | SS13 | logld2 | 0.058439 |
| 3 | ctg7180001928111_1958_SCT | 70073156 | SS13 | logld2 | -0.06564 |
| 3 | ctg7180001928111_2345_SAG | 70073543 | SS13 | logld2 | -0.06564 |
| 9 | ctg7180001848942_3677_SAG | 62019535 | SS13 | logld1 | -0.06934 |
| 9 | ctg7180001895300_2474_SAC | 64899373 | SS13 | logld1 | -0.06463 |
| 9 | ctg7180001895300_4798_SAC | 64901965 | SS13 | logld1 | -0.06463 |
| 14 | ctg7180001871200_12931_SCT | 21336770 | SS13 | logld1 | -0.0652 |
| 14 | ctg7180001871200_12984_SGT | 21336823 | SS13 | logld1 | -0.0652 |
| 14 | ctg7180001871200_12988_SAG | 21336827 | SS13 | logld1 | -0.0652 |
| 14 | ctg7180001871200_13245_SCT | 21337084 | SS13 | logld1 | -0.0652 |
| 9 | ctg7180001581419_1327_SCT | 71960485 | SS13 | logld1 | -0.07642 |
| 18 | ctg7180001791345_3710_SCT | 66814398 | SS13 | logld1 | -0.0663 |
| 18 | ctg7180001791345_3473_SAC | 66814635 | SS13 | logld1 | -0.0663 |
| 3 | ctg7180001825788_12570_SAC | 70193555 | SS13 | ld2 | -0.05729 |
| 26 | ctg7180001809454_9640_SGT | 11133973 | SS13 | ld2 | 0.049076 |
| 26 | ctg7180001809454_6285_SCT | 11137328 | SS13 | ld2 | 0.049076 |
| 26 | ctg7180001809454_6259_SAC | 11137354 | SS13 | ld2 | 0.049076 |
| 26 | ctg7180001809454_6191_SAG | 11137422 | SS13 | ld2 | 0.049076 |
| 26 | ctg7180001809454_6104_SAG | 11137509 | SS13 | ld2 | 0.049076 |
| 26 | ctg7180001809454_3127_SAC | 11140486 | SS13 | ld2 | 0.049076 |
| 26 | ctg7180001809454_2266_SAC | 11141347 | SS13 | ld2 | 0.049076 |
| 10 | ctg7180001900356_9858_SAC | 11112029 | SS13 | ld2 | 0.055703 |
| 1 | ctg7180001824066_387_SAC | 62013717 | SS13 | ld2 | -0.04139 |
| 3 | ctg7180001870985_5161_SGT | 13909130 | SS13 | ld1 | 0.092192 |
| 14 | ctg7180001879286_11274_SCT | 15466046 | SS13 | ld1 | 0.0549 |
| 14 | ctg7180001879286_13962_SAG | 15468734 | SS13 | ld1 | 0.0549 |
| 3 | ctg7180001710177_1489_SCT | 6877116 | SS13 | ld1 | -0.05158 |
| 3 | ctg7180001192621_3473_SCT | 6881980 | SS13 | ld1 | -0.05158 |
| 3 | ctg7180001710177_2453_SAG | 6878080 | SS13 | ld1 | -0.05244 |

| | | | | | |
|---|---|---|---|---|---|
| 14 | ctg7180001879286_15180_SAG | 15469952 | SS13 | ld1 | 0.054377 |
| 14 | ctg7180001879286_15189_SCT | 15469961 | SS13 | ld1 | 0.054377 |
| 3 | ctg7180001893676_9789_SCG | 5796712 | SS13 | ld1 | 0.057351 |
| 3 | ctg7180001893676_2347_SAG | 5804154 | SS13 | ld1 | 0.057351 |
| 3 | ctg7180001893676_729_SGT | 5805772 | SS13 | ld1 | 0.057351 |
| 3 | ctg7180001653065_1057_SGT | 5807756 | SS13 | ld1 | 0.057351 |
| 3 | ctg7180001653065_1850_SAC | 5808549 | SS13 | ld1 | 0.057351 |
| 4 | ctg7180001895131_7274_SCT | 65578770 | SS13 | lc2 | 3.14085 |
| 9 | ctg7180001361659_1650_SAG | 6064750 | SS13 | lc2 | 1.45449 |
| 4 | ctg7180001489651_2145_SGT | 61558925 | SS13 | lc2 | 2.91508 |
| 4 | ctg7180001898179_32187_SCT | 65079867 | SS13 | lc2 | 1.97617 |
| 4 | ctg7180001898179_32938_SAC | 65080618 | SS13 | lc2 | 1.97617 |
| 4 | ctg7180001685680_13212_SCT | 65099091 | SS13 | lc2 | 1.97617 |
| 4 | ctg7180001685680_12753_SAC | 65099550 | SS13 | lc2 | 1.97617 |
| 4 | ctg7180001685680_2479_SGT | 65109824 | SS13 | lc2 | 1.97617 |
| 4 | ctg7180001825936_14494_SAC | 50402233 | SS13 | lc2 | 1.5515 |
| 4 | ctg7180001937212_7800_SAC | 60153329 | SS13 | lc2 | 2.78152 |
| 4 | ctg7180001937212_11553_SAG | 60157082 | SS13 | lc2 | 2.78152 |
| 3 | ctg7180001710177_2453_SAG | 6878080 | SS13 | lc1 | -1.65574 |
| 3 | ctg7180001710177_1489_SCT | 6877116 | SS13 | lc1 | -1.62257 |
| 3 | ctg7180001192621_3473_SCT | 6881980 | SS13 | lc1 | -1.62257 |
| 15 | ctg7180001849195_1182_SAG | 71245010 | SS13 | lc1 | 2.96639 |
| 3 | ctg7180001893676_9789_SCG | 5796712 | SS13 | lc1 | 1.79876 |
| 3 | ctg7180001893676_2347_SAG | 5804154 | SS13 | lc1 | 1.79876 |
| 3 | ctg7180001893676_729_SGT | 5805772 | SS13 | lc1 | 1.79876 |
| 3 | ctg7180001653065_1057_SGT | 5807756 | SS13 | lc1 | 1.79876 |
| 3 | ctg7180001653065_1850_SAC | 5808549 | SS13 | lc1 | 1.79876 |
| 9 | ctg7180001900246_2441_SAG | 62406775 | SS13 | lc1 | -1.87316 |
| 20 | ctg7180001816269_19373_SAG | 46296074 | SS16 | logld2 | 0.169963 |
| 6 | ctg7180001863804_24871_SAG | 57196073 | SS16 | logld2 | -0.13626 |
| 20 | ctg7180001872068_19381_SAG | 46333593 | SS16 | logld2 | 0.167754 |
| 15 | ctg7180001910818_17819_SAG | 59951458 | SS16 | logld2 | 0.08994 |
| 15 | ctg7180001910818_18057_SAC | 59951696 | SS16 | logld2 | 0.08994 |
| 15 | ctg7180001910818_3483_SAC | 59937122 | SS16 | logld2 | 0.088828 |
| 15 | ctg7180001910818_18345_SCT | 59951984 | SS16 | logld2 | 0.088733 |
| 19 | ctg7180001717761_2022_SAG | 47364290 | SS16 | logld2 | 0.095623 |
| 14 | ctg7180001426167_1324_SGT | 17628393 | SS16 | logld2 | -0.08491 |
| 6 | ctg7180001889364_880_SAG | 60596027 | SS16 | logld2 | -0.11746 |
| 18 | ctg7180001930555_3629_SAG | 35916756 | SS16 | logld1 | -0.12212 |
| 18 | ctg7180001930555_7012_SCT | 35920139 | SS16 | logld1 | -0.12212 |
| 18 | ctg7180001930555_11273_SGT | 35924400 | SS16 | logld1 | -0.12212 |
| 18 | ctg7180001930555_12312_SAC | 35925439 | SS16 | logld1 | -0.12212 |
| 18 | ctg7180001840817_3499_SGT | 35929883 | SS16 | logld1 | -0.12212 |
| 9 | ctg7180001896160_14167_SCT | 15340857 | SS16 | logld1 | 0.092443 |
| 9 | ctg7180001642829_7752_SAG | 14708066 | SS16 | logld1 | 0.094701 |
| 21 | ctg7180001904341_12929_SAG | 19160681 | SS16 | logld1 | -0.09411 |
| 26 | ctg7180001900123_954_SGT | 37947320 | SS16 | logld1 | -0.08897 |
| 21 | ctg7180001568998_153_SCT | 19739520 | SS16 | logld1 | -0.0962 |
| 21 | ctg7180001568998_132_SGT | 19739541 | SS16 | logld1 | -0.0962 |
| 20 | ctg7180001816269_19373_SAG | 46296074 | SS16 | ld2 | 0.159167 |

| | | | | | |
|---|---|---|---|---|---|
| 20 | ctg7180001872068_19381_SAG | 46333593 | SS16 | ld2 | 0.158188 |
| 20 | ctg7180001816269_24556_SCT | 46301257 | SS16 | ld2 | 0.147892 |
| 20 | ctg7180001816269_25953_SCT | 46302654 | SS16 | ld2 | 0.147892 |
| 12 | ctg7180001880090_10977_SGT | 35642433 | SS16 | ld2 | 0.152962 |
| 6 | ctg7180001888567_3401_SCT | 42325532 | SS16 | ld2 | -0.1358 |
| 21 | ctg7180001930581_9941_SCT | 25089001 | SS16 | ld2 | 0.123116 |
| 16 | ctg7180001643134_2230_SCT | 33793228 | SS16 | ld2 | -0.10445 |
| 12 | ctg7180001876033_1027_SAC | 35609883 | SS16 | ld2 | 0.147982 |
| 5 | ctg7180001937612_17441_SAG | 41317239 | SS16 | ld2 | 0.16261 |
| 26 | ctg7180001805366_1490_SCT | 23220959 | SS16 | ld1 | 0.132222 |
| 26 | ctg7180001356609_7346_SAG | 23288731 | SS16 | ld1 | 0.14281 |
| 26 | ctg7180001878749_8526_SGT | 23336718 | SS16 | ld1 | 0.140136 |
| 26 | ctg7180001878749_11325_SCT | 23339517 | SS16 | ld1 | 0.140136 |
| 26 | ctg7180001878749_12768_SAC | 23340960 | SS16 | ld1 | 0.140136 |
| 26 | ctg7180001807110_1646_SGT | 23349945 | SS16 | ld1 | 0.140136 |
| 26 | ctg7180001807110_7984_SAG | 23356283 | SS16 | ld1 | 0.140136 |
| 26 | GCR_cBin30787_Ctg1_257 | 23391523 | SS16 | ld1 | 0.139941 |
| 26 | ctg7180001618691_1236_SAG | 21297275 | SS16 | ld1 | 0.130498 |
| 26 | ctg7180001618691_908_SCT | 21297603 | SS16 | ld1 | 0.130498 |
| 12 | ctg7180001880090_10977_SGT | 35642433 | SS16 | lc2 | 5.67922 |
| 21 | ctg7180001887190_19366_SGT | 45210909 | SS16 | lc2 | 6.35825 |
| 21 | ctg7180001906603_5378_SAC | 45247948 | SS16 | lc2 | 6.35825 |
| 21 | ctg7180001909539_5145_SCT | 45346501 | SS16 | lc2 | 6.35825 |
| 21 | ctg7180001860747_23224_SAC | 45447105 | SS16 | lc2 | 6.35825 |
| 21 | ctg7180001860747_11807_SGT | 45458522 | SS16 | lc2 | 6.35825 |
| 21 | ctg7180001930581_9941_SCT | 25089001 | SS16 | lc2 | 4.52088 |
| 20 | ctg7180001816269_19373_SAG | 46296074 | SS16 | lc2 | 5.32282 |
| 12 | ctg7180001876033_1027_SAC | 35609883 | SS16 | lc2 | 5.49267 |
| 20 | ctg7180001872068_19381_SAG | 46333593 | SS16 | lc2 | 5.30857 |
| 26 | ctg7180001805366_1490_SCT | 23220959 | SS16 | lc1 | 3.12843 |
| 21 | ctg7180001568998_153_SCT | 19739520 | SS16 | lc1 | -3.44014 |
| 21 | ctg7180001568998_132_SGT | 19739541 | SS16 | lc1 | -3.44014 |
| 21 | ctg7180001580620_1390_SAG | 19728394 | SS16 | lc1 | -3.43778 |
| 26 | ctg7180001356609_7346_SAG | 23288731 | SS16 | lc1 | 3.38195 |
| 21 | ctg7180001904341_12929_SAG | 19160681 | SS16 | lc1 | -3.25317 |
| 26 | GCR_cBin30787_Ctg1_257 | 23391523 | SS16 | lc1 | 3.31517 |
| 26 | ctg7180001878749_8526_SGT | 23336718 | SS16 | lc1 | 3.30739 |
| 26 | ctg7180001878749_11325_SCT | 23339517 | SS16 | lc1 | 3.30739 |
| 26 | ctg7180001878749_12768_SAC | 23340960 | SS16 | lc1 | 3.30739 |
| 26 | ctg7180001807110_1646_SGT | 23349945 | SS16 | lc1 | 3.30739 |
| 26 | ctg7180001807110_7984_SAG | 23356283 | SS16 | lc1 | 3.30739 |
| 3 | ctg7180001865965_4471_SCT | 78352498 | SS17 | logld4 | 0.087341 |
| 3 | ctg7180001865965_4181_SAG | 78352788 | SS17 | logld4 | 0.087341 |
| 12 | ctg7180001544960_4389_SCT | 80810648 | SS17 | logld4 | -0.10936 |
| 12 | AX-87821834 | 76998041 | SS17 | logld4 | -0.11011 |
| 12 | ctg7180001516140_2173_SAC | 79410169 | SS17 | logld4 | -0.12165 |
| 12 | ctg7180001832513_16579_SCT | 75220021 | SS17 | logld4 | -0.07965 |
| 10 | ctg7180001229400_1707_SGT | 67921632 | SS17 | logld4 | -0.07487 |
| 12 | ctg7180001544960_3033_SCT | 80809138 | SS17 | logld4 | -0.11824 |
| 3 | ctg7180001803004_1580_SAG | 79209105 | SS17 | logld4 | 0.093437 |

| | | | | | |
|---|---|---|---|---|---|
| 3 | ctg7180001460681_1054_SAG | 78223769 | SS17 | logld4 | 0.079348 |
| 12 | ctg7180001774501_774_SAG | 75519499 | SS17 | logld3 | 0.130615 |
| 12 | ctg7180001845361_4514_SAC | 14653588 | SS17 | logld3 | -0.1219 |
| 2 | ctg7180001812661_8524_SCT | 31869150 | SS17 | logld3 | -0.11185 |
| 12 | ctg7180001885480_4483_SAC | 14664002 | SS17 | logld3 | -0.08812 |
| 2 | ctg7180001812661_13610_SCT | 31864064 | SS17 | logld3 | -0.11185 |
| 2 | ctg7180001917965_14405_SGT | 31861535 | SS17 | logld3 | -0.11865 |
| 12 | ctg7180001885480_1793_SCT | 14661312 | SS17 | logld3 | -0.11911 |
| 12 | ctg7180001845361_5456_SGT | 14652646 | SS17 | logld3 | -0.11818 |
| 8 | ctg7180001518605_1040_SGT | 11817897 | SS17 | logld3 | -0.08387 |
| 15 | ctg7180001839857_9373_SGT | 22955947 | SS17 | logld3 | -0.08324 |
| 15 | ctg7180001839857_9384_SAT | 22955958 | SS17 | logld3 | -0.08324 |
| 12 | AX-96443306 | 15637088 | SS17 | logld2 | -0.13761 |
| 12 | ctg7180001883118_1758_SAG | 15626305 | SS17 | logld2 | -0.13248 |
| 12 | ctg7180001838384_1351_SAC | 14045429 | SS17 | logld2 | 0.1171 |
| 12 | AX-87074389 | 15608796 | SS17 | logld2 | -0.12592 |
| 1 | ctg7180001801278_2651_SAC | 128368389 | SS17 | logld2 | -0.22161 |
| 22 | ctg7180001911652_10325_SCT | 21112207 | SS17 | logld2 | 0.1261 |
| 1 | ctg7180001924116_1476_SAG | 128364076 | SS17 | logld2 | -0.21605 |
| 1 | ctg7180001924116_438_SCT | 128365114 | SS17 | logld2 | -0.21605 |
| 1 | ctg7180001924116_400_SAC | 128365152 | SS17 | logld2 | -0.21605 |
| 22 | ctg7180001911652_7656_SCT | 21109538 | SS17 | logld2 | 0.125449 |
| 14 | ctg7180001514052_409_SAC | 86111056 | SS17 | logld1 | -0.23596 |
| 14 | ctg7180001514052_717_SAG | 86111267 | SS17 | logld1 | -0.23596 |
| 14 | ctg7180001514052_936_SAG | 86111486 | SS17 | logld1 | -0.23596 |
| 19 | ctg7180001903927_3991_SAG | 77911367 | SS17 | logld1 | 0.165058 |
| 17 | ctg7180001726890_1710_SGT | 46354454 | SS17 | logld1 | -0.14362 |
| 14 | ctg7180001847610_15837_SCG | 8739295 | SS17 | logld1 | 0.136768 |
| 14 | ctg7180001847610_16289_SCT | 8739747 | SS17 | logld1 | 0.136768 |
| 15 | ctg7180001310117_8573_SAC | 25925374 | SS17 | logld1 | -0.22611 |
| 17 | ctg7180001935834_4289_SAC | 34595429 | SS17 | logld1 | -0.14095 |
| 17 | ctg7180001894313_6896_SAC | 54018754 | SS17 | logld1 | 0.192744 |
| 3 | ctg7180001865965_4471_SCT | 78352498 | SS17 | ld4 | 0.048856 |
| 3 | ctg7180001865965_4181_SAG | 78352788 | SS17 | ld4 | 0.048856 |
| 3 | ctg7180001460681_1054_SAG | 78223769 | SS17 | ld4 | 0.046554 |
| 3 | ctg7180001268912_1361_SGT | 62424813 | SS17 | ld4 | 0.051578 |
| 3 | ctg7180001846161_2434_SAG | 62444658 | SS17 | ld4 | 0.051218 |
| 3 | ctg7180001865611_5234_SCT | 76277157 | SS17 | ld4 | 0.046391 |
| 3 | ctg7180001927258_6212_SCT | 76157704 | SS17 | ld4 | 0.046915 |
| 3 | ctg7180001645083_2635_SGT | 76273429 | SS17 | ld4 | 0.045923 |
| 3 | ctg7180001645083_1493_SGT | 76274571 | SS17 | ld4 | 0.04581 |
| 3 | ctg7180001645083_1286_SCT | 76274778 | SS17 | ld4 | 0.04581 |
| 3 | ctg7180001865965_4471_SCT | 78352498 | SS17 | ld3 | 0.030259 |
| 3 | ctg7180001865965_4181_SAG | 78352788 | SS17 | ld3 | 0.030259 |
| 12 | ctg7180001774501_774_SAG | 75519499 | SS17 | ld3 | 0.043906 |
| 12 | ctg7180001885480_4483_SAC | 14664002 | SS17 | ld3 | -0.03012 |
| 12 | ctg7180001845361_4514_SAC | 14653588 | SS17 | ld3 | -0.03994 |
| 17 | ctg7180001815508_106_SGT | 30331957 | SS17 | ld3 | 0.04052 |
| 17 | ctg7180001801849_1431_SAG | 28165439 | SS17 | ld3 | 0.040591 |
| 3 | AX-87468197 | 77320232 | SS17 | ld3 | 0.028843 |

| | | | | | |
|---|---|---|---|---|---|
| 3 | ctg7180001702615_1060_SCT | 56214045 | SS17 | ld3 | 0.032381 |
| 17 | ctg7180001817400_6576_SAG | 30326877 | SS17 | ld3 | 0.039784 |
| 12 | AX-96443306 | 15637088 | SS17 | ld2 | -0.05605 |
| 12 | AX-87074389 | 15608796 | SS17 | ld2 | -0.05462 |
| 12 | ctg7180001883118_1892_SGT | 15626439 | SS17 | ld2 | -0.05254 |
| 12 | ctg7180001883118_1758_SAG | 15626305 | SS17 | ld2 | -0.05156 |
| 12 | ctg7180001838384_1351_SAC | 14045429 | SS17 | ld2 | 0.045529 |
| 12 | ctg7180001578334_2039_SAC | 13656675 | SS17 | ld2 | 0.045958 |
| 12 | ctg7180001181899_1233_SGT | 14612392 | SS17 | ld2 | -0.05128 |
| 12 | ctg7180001181899_1001_SCT | 14612624 | SS17 | ld2 | -0.05128 |
| 12 | ctg7180001181899_1266_SAC | 14612359 | SS17 | ld2 | -0.05095 |
| 12 | ctg7180001935730_27198_SCT | 13531143 | SS17 | ld2 | -0.0473 |
| 17 | ctg7180001894313_6896_SAC | 54018754 | SS17 | ld1 | 0.049066 |
| 11 | ctg7180001937375_45126_SAC | 58913719 | SS17 | ld1 | 0.032754 |
| 13 | AX-87313021 | 52392573 | SS17 | ld1 | 0.035529 |
| 17 | ctg7180001822016_322_SGT | 31405928 | SS17 | ld1 | 0.063362 |
| 17 | ctg7180001835615_1880_SAG | 31549935 | SS17 | ld1 | 0.063362 |
| 17 | ctg7180001884700_454_SAG | 31584705 | SS17 | ld1 | 0.063362 |
| 13 | ctg7180001933044_11868_SAC | 52231201 | SS17 | ld1 | 0.035863 |
| 17 | ctg7180001822781_4585_SCT | 44855802 | SS17 | ld1 | 0.029415 |
| 17 | ctg7180001822781_4952_SAG | 44856168 | SS17 | ld1 | 0.029415 |
| 17 | ctg7180001822781_6489_SAG | 44857705 | SS17 | ld1 | 0.029415 |
| 17 | ctg7180001822781_10297_SCT | 44861513 | SS17 | ld1 | 0.029415 |
| 3 | ctg7180001268912_1361_SGT | 62424813 | SS17 | lc4 | 2.97055 |
| 3 | ctg7180001846161_2434_SAG | 62444658 | SS17 | lc4 | 2.95794 |
| 3 | ctg7180001460681_1054_SAG | 78223769 | SS17 | lc4 | 2.62706 |
| 3 | AX-215398188 | 62423318 | SS17 | lc4 | 2.9409 |
| 3 | ctg7180001920538_9002_SGT | 62493347 | SS17 | lc4 | 2.84286 |
| 3 | ctg7180001799042_5615_SCT | 76854747 | SS17 | lc4 | 3.39095 |
| 3 | ctg7180001883197_436_SCT | 76918934 | SS17 | lc4 | 3.39095 |
| 3 | ctg7180001796000_1835_SAT | 74845335 | SS17 | lc4 | 3.40888 |
| 3 | ctg7180001796000_4990_SCT | 74850282 | SS17 | lc4 | 3.40888 |
| 3 | ctg7180001796001_3928_SAG | 74859032 | SS17 | lc4 | 3.40888 |
| 12 | ctg7180001774501_774_SAG | 75519499 | SS17 | lc3 | 2.5388 |
| 17 | ctg7180001801849_1431_SAG | 28165439 | SS17 | lc3 | 2.46056 |
| 17 | ctg7180001815508_106_SGT | 30331957 | SS17 | lc3 | 2.36725 |
| 3 | ctg7180001702615_1060_SCT | 56214045 | SS17 | lc3 | 1.91086 |
| 17 | ctg7180001817400_6576_SAG | 30326877 | SS17 | lc3 | 2.32769 |
| 12 | ctg7180001427127_551_SCT | 69100338 | SS17 | lc3 | 2.80573 |
| 13 | ctg7180001831038_2520_SAG | 11746508 | SS17 | lc3 | 3.32319 |
| 12 | AX-87060120 | 34628507 | SS17 | lc3 | -1.56763 |
| 17 | ctg7180001721010_67_SCT | 33278393 | SS17 | lc3 | 3.04586 |
| 12 | ctg7180001845361_4514_SAC | 14653588 | SS17 | lc3 | -2.26406 |
| 12 | AX-96443306 | 15637088 | SS17 | lc2 | -2.21294 |
| 12 | AX-87074389 | 15608796 | SS17 | lc2 | -2.11977 |
| 12 | ctg7180001883118_1892_SGT | 15626439 | SS17 | lc2 | -2.05153 |
| 12 | ctg7180001883118_1758_SAG | 15626305 | SS17 | lc2 | -2.06637 |
| 2 | ctg7180001924737_4228_SAG | 55924783 | SS17 | lc2 | 1.91564 |
| 12 | ctg7180001889505_11144_SAC | 13416450 | SS17 | lc2 | 2.05414 |
| 3 | ctg7180001796000_1835_SAT | 74845335 | SS17 | lc2 | 2.47385 |

| | | | | | |
|---|---|---|---|---|---|
| 3 | ctg7180001796000_4990_SCT | 74850282 | SS17 | lc2 | 2.47385 |
| 3 | ctg7180001796001_3928_SAG | 74859032 | SS17 | lc2 | 2.47385 |
| 12 | ctg7180001892628_7267_SGT | 20180730 | SS17 | lc2 | 1.86773 |
| 11 | ctg7180001937375_45126_SAC | 58913719 | SS17 | lc1 | 1.41856 |
| 17 | ctg7180001894313_6896_SAC | 54018754 | SS17 | lc1 | 1.98646 |
| 17 | ctg7180001746484_4788_SAG | 53859082 | SS17 | lc1 | 2.35702 |
| 11 | AX-87323716 | 58923059 | SS17 | lc1 | 1.31916 |
| 17 | ctg7180001822016_322_SGT | 31405928 | SS17 | lc1 | 2.59863 |
| 17 | ctg7180001835615_1880_SAG | 31549935 | SS17 | lc1 | 2.59863 |
| 17 | ctg7180001884700_454_SAG | 31584705 | SS17 | lc1 | 2.59863 |
| 17 | ctg7180001822781_4585_SCT | 44855802 | SS17 | lc1 | 1.20857 |
| 17 | ctg7180001822781_4952_SAG | 44856168 | SS17 | lc1 | 1.20857 |
| 17 | ctg7180001822781_6489_SAG | 44857705 | SS17 | lc1 | 1.20857 |
| 17 | ctg7180001822781_10297_SCT | 44861513 | SS17 | lc1 | 1.20857 |

| MAF | p-value | Variance_explained |
|---|---|---|
| 0.1706 | 1.24E-08 | 0.00613156 |
| 0.1647 | 2.00E-06 | 0.003934771 |
| 0.1647 | 2.00E-06 | 0.003934771 |
| 0.1647 | 2.00E-06 | 0.003934771 |
| 0.111 | 6.71E-06 | 0.003643466 |
| 0.1428 | 7.38E-06 | 0.002931818 |
| 0.1388 | 8.46E-06 | 0.003744425 |
| 0.3703 | 8.56E-06 | 0.003189867 |
| 0.2403 | 8.89E-06 | 0.00343169 |
| 0.4375 | 9.04E-06 | 0.003676295 |
| 0.4865 | 8.77E-08 | 0.003990846 |
| 0.4737 | 1.51E-07 | 0.003805353 |
| 0.4737 | 1.51E-07 | 0.003805353 |
| 0.4737 | 1.51E-07 | 0.003805353 |
| 0.4953 | 4.34E-07 | 0.00351968 |
| 0.4953 | 4.34E-07 | 0.00351968 |
| 0.3408 | 1.87E-06 | 0.002988903 |
| 0.3411 | 2.37E-06 | 0.002944398 |
| 0.4478 | 4.88E-06 | 0.002727735 |
| 0.1399 | 5.27E-06 | 0.00309817 |
| 0.1706 | 7.52E-09 | 0.002680482 |
| 0.1388 | 3.60E-07 | 0.00207495 |
| 0.4375 | 3.90E-07 | 0.002034204 |
| 0.1647 | 7.46E-07 | 0.001801908 |
| 0.1647 | 7.46E-07 | 0.001801908 |
| 0.1647 | 7.46E-07 | 0.001801908 |
| 0.1428 | 1.04E-06 | 0.001458907 |
| 0.05917 | 1.20E-06 | 0.00175617 |
| 0.2425 | 1.21E-06 | 0.001798492 |
| 0.1384 | 1.41E-06 | 0.001859769 |
| 0.1384 | 1.41E-06 | 0.001859769 |
| 0.4865 | 7.32E-08 | 0.003684089 |
| 0.4737 | 1.34E-07 | 0.003500001 |
| 0.4737 | 1.34E-07 | 0.003500001 |
| 0.4737 | 1.34E-07 | 0.003500001 |
| 0.4953 | 5.17E-07 | 0.00316484 |
| 0.4953 | 5.17E-07 | 0.00316484 |
| 0.3408 | 1.89E-06 | 0.002720326 |
| 0.3411 | 2.33E-06 | 0.002685819 |
| 0.4134 | 4.01E-06 | 0.002776055 |
| 0.351 | 5.18E-06 | 0.002562534 |
| 0.1388 | 1.25E-07 | 2.699798069 |
| 0.08875 | 3.12E-07 | 2.36776966 |
| 0.1384 | 4.88E-07 | 2.441215632 |
| 0.1384 | 4.88E-07 | 2.441215632 |
| 0.1545 | 4.98E-07 | 2.510412238 |
| 0.1545 | 4.98E-07 | 2.510412238 |
| 0.1804 | 5.21E-07 | 2.402305814 |
| 0.1804 | 5.21E-07 | 2.402305814 |

| | | |
|---|---|---|
| 0.1644 | 1.10E-06 | 2.265305123 |
| 0.1538 | 1.18E-06 | 2.321661674 |
| 0.4865 | 3.58E-07 | 2.898875285 |
| 0.4737 | 6.99E-07 | 2.722856202 |
| 0.4737 | 6.99E-07 | 2.722856202 |
| 0.4737 | 6.99E-07 | 2.722856202 |
| 0.4953 | 2.66E-06 | 2.468290564 |
| 0.4953 | 2.66E-06 | 2.468290564 |
| 0.351 | 8.12E-06 | 2.206156903 |
| 0.3305 | 1.08E-05 | 2.243563565 |
| 0.3305 | 1.08E-05 | 2.243563565 |
| 0.3444 | 1.11E-05 | 2.377701971 |
| 0.3444 | 1.11E-05 | 2.377701971 |
| 0.175 | 5.12E-07 | 0.00172155 |
| 0.3307 | 1.37E-06 | 0.001827794 |
| 0.3307 | 1.37E-06 | 0.001827794 |
| 0.3307 | 1.37E-06 | 0.001827794 |
| 0.3307 | 1.37E-06 | 0.001827794 |
| 0.3307 | 1.37E-06 | 0.001827794 |
| 0.3307 | 1.37E-06 | 0.001827794 |
| 0.3307 | 1.37E-06 | 0.001827794 |
| 0.3102 | 2.45E-06 | 0.001461481 |
| 0.2313 | 2.62E-06 | 0.001532294 |
| 0.2313 | 2.62E-06 | 0.001532294 |
| 0.3341 | 5.61E-07 | 0.00213911 |
| 0.4073 | 8.55E-07 | 0.002016511 |
| 0.4073 | 8.55E-07 | 0.002016511 |
| 0.2892 | 1.13E-06 | 0.001747979 |
| 0.2892 | 1.13E-06 | 0.001747979 |
| 0.2892 | 1.13E-06 | 0.001747979 |
| 0.2892 | 1.13E-06 | 0.001747979 |
| 0.2661 | 1.14E-06 | 0.002280842 |
| 0.4174 | 1.14E-06 | 0.002137606 |
| 0.4174 | 1.14E-06 | 0.002137606 |
| 0.175 | 9.41E-07 | 0.000947789 |
| 0.3307 | 1.24E-06 | 0.001066141 |
| 0.3307 | 1.24E-06 | 0.001066141 |
| 0.3307 | 1.24E-06 | 0.001066141 |
| 0.3307 | 1.24E-06 | 0.001066141 |
| 0.3307 | 1.24E-06 | 0.001066141 |
| 0.3307 | 1.24E-06 | 0.001066141 |
| 0.3307 | 1.24E-06 | 0.001066141 |
| 0.223 | 1.36E-06 | 0.00107524 |
| 0.4634 | 1.85E-06 | 0.000851886 |
| 0.08333 | 3.42E-07 | 0.001298458 |
| 0.4478 | 6.52E-07 | 0.001490601 |
| 0.4478 | 6.52E-07 | 0.001490601 |
| 0.3884 | 8.01E-07 | 0.001263938 |
| 0.3884 | 8.01E-07 | 0.001263938 |
| 0.3988 | 8.12E-07 | 0.001318439 |

| | | |
|---|---|---|
| 0.4483 | 8.14E-07 | 0.001462606 |
| 0.4483 | 8.14E-07 | 0.001462606 |
| 0.2656 | 1.24E-06 | 0.001283141 |
| 0.2656 | 1.24E-06 | 0.001283141 |
| 0.2656 | 1.24E-06 | 0.001283141 |
| 0.2656 | 1.24E-06 | 0.001283141 |
| 0.2656 | 1.24E-06 | 0.001283141 |
| 0.05659 | 2.39E-06 | 1.053330251 |
| 0.3102 | 2.64E-06 | 0.905349901 |
| 0.05919 | 7.86E-06 | 0.946414131 |
| 0.1163 | 1.04E-05 | 0.80271831 |
| 0.1163 | 1.04E-05 | 0.80271831 |
| 0.1163 | 1.04E-05 | 0.80271831 |
| 0.1163 | 1.04E-05 | 0.80271831 |
| 0.1163 | 1.04E-05 | 0.80271831 |
| 0.2144 | 1.14E-05 | 0.810886017 |
| 0.06205 | 1.34E-05 | 0.900566615 |
| 0.06205 | 1.34E-05 | 0.900566615 |
| 0.3988 | 5.03E-07 | 1.314584171 |
| 0.3884 | 5.44E-07 | 1.25078763 |
| 0.3884 | 5.44E-07 | 1.25078763 |
| 0.06983 | 9.98E-07 | 1.143117473 |
| 0.2656 | 1.04E-06 | 1.262226161 |
| 0.2656 | 1.04E-06 | 1.262226161 |
| 0.2656 | 1.04E-06 | 1.262226161 |
| 0.2656 | 1.04E-06 | 1.262226161 |
| 0.2656 | 1.04E-06 | 1.262226161 |
| 0.2131 | 1.66E-06 | 1.176746028 |
| 0.07837 | 2.70E-06 | 0.00417297 |
| 0.1872 | 3.83E-06 | 0.005649853 |
| 0.07771 | 3.94E-06 | 0.004033854 |
| 0.4253 | 4.91E-06 | 0.003954334 |
| 0.4253 | 4.91E-06 | 0.003954334 |
| 0.4312 | 5.29E-06 | 0.003870474 |
| 0.4263 | 6.39E-06 | 0.003851205 |
| 0.46 | 7.09E-06 | 0.004542619 |
| 0.3783 | 7.17E-06 | 0.003391002 |
| 0.2189 | 7.90E-06 | 0.004717807 |
| 0.1663 | 1.27E-06 | 0.004135558 |
| 0.1663 | 1.27E-06 | 0.004135558 |
| 0.1663 | 1.27E-06 | 0.004135558 |
| 0.1663 | 1.27E-06 | 0.004135558 |
| 0.1663 | 1.27E-06 | 0.004135558 |
| 0.4296 | 2.08E-06 | 0.004188137 |
| 0.3009 | 3.94E-06 | 0.003773104 |
| 0.2421 | 4.18E-06 | 0.003250297 |
| 0.3776 | 4.67E-06 | 0.003721034 |
| 0.2159 | 5.33E-06 | 0.003133473 |
| 0.2159 | 5.33E-06 | 0.003133473 |
| 0.07837 | 4.57E-07 | 0.003659675 |

| | | |
|---|---|---|
| 0.07771 | 5.89E-07 | 0.003586918 |
| 0.08234 | 1.50E-06 | 0.003305309 |
| 0.08234 | 1.50E-06 | 0.003305309 |
| 0.07308 | 3.34E-06 | 0.003169844 |
| 0.08962 | 3.34E-06 | 0.003009375 |
| 0.1118 | 3.75E-06 | 0.003010312 |
| 0.2014 | 5.47E-06 | 0.003509624 |
| 0.07341 | 5.89E-06 | 0.002979138 |
| 0.06151 | 6.03E-06 | 0.003052811 |
| 0.3366 | 7.78E-07 | 0.00780777 |
| 0.251 | 9.98E-07 | 0.007668365 |
| 0.25 | 1.60E-06 | 0.007364287 |
| 0.25 | 1.60E-06 | 0.007364287 |
| 0.25 | 1.60E-06 | 0.007364287 |
| 0.25 | 1.60E-06 | 0.007364287 |
| 0.25 | 1.60E-06 | 0.007364287 |
| 0.2512 | 1.64E-06 | 0.00736725 |
| 0.3224 | 1.65E-06 | 0.007440569 |
| 0.3224 | 1.65E-06 | 0.007440569 |
| 0.07308 | 1.16E-06 | 4.369665296 |
| 0.0506 | 1.58E-06 | 3.884230014 |
| 0.0506 | 1.58E-06 | 3.884230014 |
| 0.0506 | 1.58E-06 | 3.884230014 |
| 0.0506 | 1.58E-06 | 3.884230014 |
| 0.0506 | 1.58E-06 | 3.884230014 |
| 0.1118 | 1.71E-06 | 4.059088563 |
| 0.07837 | 2.02E-06 | 4.092795125 |
| 0.07341 | 2.16E-06 | 4.104307047 |
| 0.07771 | 2.34E-06 | 4.039517569 |
| 0.3366 | 1.67E-06 | 4.370915979 |
| 0.2159 | 1.68E-06 | 4.006879905 |
| 0.2159 | 1.68E-06 | 4.006879905 |
| 0.2169 | 1.71E-06 | 4.014790917 |
| 0.251 | 2.07E-06 | 4.300509387 |
| 0.2421 | 2.80E-06 | 3.88374055 |
| 0.2512 | 3.28E-06 | 4.134538819 |
| 0.25 | 3.47E-06 | 4.10206073 |
| 0.25 | 3.47E-06 | 4.10206073 |
| 0.25 | 3.47E-06 | 4.10206073 |
| 0.25 | 3.47E-06 | 4.10206073 |
| 0.25 | 3.47E-06 | 4.10206073 |
| 0.3478 | 9.55E-07 | 0.003460801 |
| 0.3478 | 9.55E-07 | 0.003460801 |
| 0.1548 | 2.57E-06 | 0.003129232 |
| 0.1577 | 2.95E-06 | 0.003220877 |
| 0.1245 | 3.07E-06 | 0.003226061 |
| 0.356 | 3.69E-06 | 0.002908929 |
| 0.4185 | 4.23E-06 | 0.002728518 |
| 0.1294 | 4.39E-06 | 0.003150062 |
| 0.2236 | 5.22E-06 | 0.003031299 |

| | | |
|---|---|---|
| 0.4155 | 5.47E-06 | 0.003058156 |
| 0.1114 | 1.29E-06 | 0.003377595 |
| 0.1189 | 2.74E-06 | 0.003113519 |
| 0.1391 | 2.84E-06 | 0.00299649 |
| 0.2771 | 2.95E-06 | 0.00311087 |
| 0.138 | 3.18E-06 | 0.00297638 |
| 0.1234 | 3.63E-06 | 0.003045408 |
| 0.1189 | 4.62E-06 | 0.002972729 |
| 0.1197 | 4.78E-06 | 0.002943157 |
| 0.2644 | 6.31E-06 | 0.002735996 |
| 0.2786 | 7.54E-06 | 0.002785014 |
| 0.2786 | 7.54E-06 | 0.002785014 |
| 0.293 | 2.30E-07 | 0.007845777 |
| 0.2801 | 8.86E-07 | 0.007078087 |
| 0.3968 | 1.33E-06 | 0.006564124 |
| 0.4695 | 2.35E-06 | 0.007898423 |
| 0.07292 | 3.66E-06 | 0.006640069 |
| 0.2599 | 4.27E-06 | 0.006117259 |
| 0.0733 | 6.15E-06 | 0.006341173 |
| 0.0733 | 6.15E-06 | 0.006341173 |
| 0.0733 | 6.15E-06 | 0.006341173 |
| 0.2524 | 7.26E-06 | 0.005939133 |
| 0.08751 | 1.25E-06 | 0.008891482 |
| 0.08751 | 1.25E-06 | 0.008891482 |
| 0.08751 | 1.25E-06 | 0.008891482 |
| 0.1829 | 3.51E-06 | 0.008143143 |
| 0.2461 | 5.52E-06 | 0.007653943 |
| 0.4817 | 5.85E-06 | 0.009340214 |
| 0.4817 | 5.85E-06 | 0.009340214 |
| 0.0531 | 1.19E-05 | 0.005141334 |
| 0.2779 | 1.23E-05 | 0.007973333 |
| 0.09461 | 1.29E-05 | 0.006364503 |
| 0.3478 | 1.12E-07 | 0.00108287 |
| 0.3478 | 1.12E-07 | 0.00108287 |
| 0.4155 | 2.45E-07 | 0.001052665 |
| 0.2659 | 3.65E-07 | 0.001038566 |
| 0.2663 | 4.05E-07 | 0.001025113 |
| 0.491 | 5.36E-07 | 0.0010757 |
| 0.3624 | 5.97E-07 | 0.00101717 |
| 0.4914 | 6.41E-07 | 0.00105414 |
| 0.4918 | 7.12E-07 | 0.00104901 |
| 0.4918 | 7.12E-07 | 0.00104901 |
| 0.3478 | 5.66E-07 | 0.000415384 |
| 0.3478 | 5.66E-07 | 0.000415384 |
| 0.1114 | 1.55E-06 | 0.00038165 |
| 0.2771 | 2.28E-06 | 0.000363446 |
| 0.1189 | 5.54E-06 | 0.000334318 |
| 0.1099 | 5.71E-06 | 0.000321219 |
| 0.1073 | 7.20E-06 | 0.000315636 |
| 0.2699 | 7.76E-06 | 0.000327859 |

| | | |
|---|---|---|
| 0.1799 | 7.90E-06 | 0.00030939 |
| 0.1103 | 8.24E-06 | 0.000310645 |
| 0.293 | 4.83E-08 | 0.001301716 |
| 0.4695 | 1.18E-07 | 0.001486334 |
| 0.4495 | 2.33E-07 | 0.001366198 |
| 0.2801 | 7.30E-07 | 0.001072069 |
| 0.3968 | 1.11E-06 | 0.000992278 |
| 0.3826 | 1.43E-06 | 0.000997825 |
| 0.2461 | 1.80E-06 | 0.00097592 |
| 0.2461 | 1.80E-06 | 0.00097592 |
| 0.2464 | 2.11E-06 | 0.00096404 |
| 0.4536 | 2.89E-06 | 0.001108875 |
| 0.09461 | 1.03E-06 | 0.000412441 |
| 0.279 | 3.03E-06 | 0.000431627 |
| 0.1914 | 6.32E-06 | 0.000390732 |
| 0.05236 | 6.49E-06 | 0.00039841 |
| 0.05236 | 6.49E-06 | 0.00039841 |
| 0.05236 | 6.49E-06 | 0.00039841 |
| 0.1877 | 6.50E-06 | 0.000392205 |
| 0.3119 | 6.84E-06 | 0.000371386 |
| 0.3119 | 6.84E-06 | 0.000371386 |
| 0.3119 | 6.84E-06 | 0.000371386 |
| 0.3119 | 6.84E-06 | 0.000371386 |
| 0.2659 | 4.14E-07 | 3.444905323 |
| 0.2663 | 4.27E-07 | 3.418994498 |
| 0.4155 | 4.82E-07 | 3.352166047 |
| 0.2626 | 1.06E-06 | 3.349564657 |
| 0.2666 | 1.10E-06 | 3.160399556 |
| 0.147 | 1.31E-06 | 2.883627335 |
| 0.147 | 1.31E-06 | 2.883627335 |
| 0.1429 | 1.33E-06 | 2.846539052 |
| 0.1429 | 1.33E-06 | 2.846539052 |
| 0.1429 | 1.33E-06 | 2.846539052 |
| 0.1114 | 8.09E-07 | 1.276081683 |
| 0.1073 | 1.36E-06 | 1.159853592 |
| 0.1099 | 2.52E-06 | 1.096363932 |
| 0.1799 | 2.85E-06 | 1.077421679 |
| 0.1103 | 3.62E-06 | 1.063406671 |
| 0.07592 | 3.70E-06 | 1.104555437 |
| 0.05797 | 3.94E-06 | 1.206169589 |
| 0.4054 | 3.94E-06 | 1.184747435 |
| 0.06058 | 3.95E-06 | 1.055939271 |
| 0.1189 | 4.85E-06 | 1.074021352 |
| 0.293 | 1.93E-07 | 2.028879751 |
| 0.4695 | 6.90E-07 | 2.23835241 |
| 0.4495 | 1.08E-06 | 2.082920812 |
| 0.2801 | 1.61E-06 | 1.721993287 |
| 0.2543 | 2.04E-06 | 1.391772633 |
| 0.2468 | 4.05E-06 | 1.568720391 |
| 0.1429 | 4.26E-06 | 1.499133971 |

| | | |
|---|---|---|
| 0.1429 | 4.26E-06 | 1.499133971 |
| 0.1429 | 4.26E-06 | 1.499133971 |
| 0.3265 | 4.42E-06 | 1.534189374 |
| 0.279 | 8.40E-07 | 0.80958953 |
| 0.09461 | 1.45E-06 | 0.676024414 |
| 0.05797 | 5.58E-06 | 0.606770649 |
| 0.279 | 5.71E-06 | 0.700106987 |
| 0.05236 | 6.67E-06 | 0.670134402 |
| 0.05236 | 6.67E-06 | 0.670134402 |
| 0.05236 | 6.67E-06 | 0.670134402 |
| 0.3119 | 6.74E-06 | 0.626961031 |
| 0.3119 | 6.74E-06 | 0.626961031 |
| 0.3119 | 6.74E-06 | 0.626961031 |
| 0.3119 | 6.74E-06 | 0.626961031 |
