## Supplementary Table 2 for "Meta-analysis of GWAS for sea lice load in Atlantic salmon"

**List of candidate genes for all QTL show in meta-analysis Manhattan plot, selected by most significant p-value:**

**LC candidate genes**

| SNP | P-VALUE | CHR | POS | Protein |
|---|---|---|---|---|
| ctg7180001924116_400_SAC | 9.50E-11 | 1 | 128365152 | Transposon Tf2-1 polyprotein<br>T-box transcription factor TBX5-A-like |
| ctg7180001664070_2019_SGT | 2.93E-08 | 1 | 802443 | Attractin isoform X2 |
| AX-87153071 | 7.48E-08 | 1 | 1148341 | Homeobox EMX1-like |
| AX-87343039 | 3.89E-10 | 1 | 150826261 | Di-Ras2 GTP-binding protein |
| AX-215443812 | 2.10E-11 | 2 | 66777452 | Zinc finger BED domain-containing protein 4-like |
| ctg7180001715025_1222_SAG | 1.22E-08 | 2 | 9536635 | Transposon Tf2-1 polyprotein |
| ctg7180001887970_1091_SCT | 2.31E-08 | 2 | 9657346 | Zinc finger 629-like |
| ctg7180001799042_5615_SCT | 1.85E-22 | 3 | 76854747 | GPTase IMAP family member 8-like |
| ctg7180001199073_3918_SCT | 4.14E-11 | 3 | 7026040 | Filamentous growth regulator 23-like isoform X1<br>Mucin-16-like isoform X2<br>Involucrin-like |
| AX-87084850 | 5.87E-12 | 4 | 62570478 | Glutamate receptor |
| ctg7180001796911_633_SGT | 4.22E-11 | 5 | 49549632 | Zinc finger protein 407-like |
| AX-215422843 | 2.37E-08 | 5 | 6471903 | Interferon alpha/beta receptor |
| AX-96439890 | 6.43E-09 | 6 | 11275392 | Dynein assembly factor 5, axonemal-like |
| ctg7180001877771_4032_SCT | 2.08E-09 | 6 | 21804607 | Inter-alpha trypsin inhibitor heavy chain H3-like isoform |
| ctg7180001912336_13519_SC<br>G | 3.94E-09 | 6 | 34428543 | KN motif and ankyrin repeat domain-containing protein 2-like<br>Hemoglobin subunit beta-1-like |
| ctg7180001255964_1180_SAG | 3.57E-09 | 6 | 37348032 | Frizzled-2-like<br>Frizzled-1-like |
| AX-86933745 | 2.03E-09 | 6 | 77405961 | alpha-(1,6)-fucosyltransferase-like isoform |
| ctg7180001931975_11476_SCT | 3.83E-09 | 7 | 31829805 | RCC2 homolog |
| AX-87426530 | 4.19E-08 | 9 | 812385 | extensin-like |
| ctg7180001889104_9124_SGT | 7.25E-15 | 9 | 56363260 | RCC2 homolog |
| ctg7180001815146_2418_SCT | 1.98E-12 | 9 | 110157594 | Calpain-5-like isoform X2 |
| AX-96495871 | 9.96E-15 | 10 | 66025311 | ZK1236.4<br>alpha-ketoglutarate-dependent dioxygenase alkB homolog 3 |
| AX-87046611 | 2.00E-13 | 10 | 2698859 | Adhesion G protein-coupled receptor<br>latrophilin-3-like isoform X11 |
| AX-87323716 | 1.89E-15 | 11 | 58923059 | microtubule-associated serine/threonine-protein kinase<br>Tf2-1 polyprotein |

| | | | | |
|---|---|---|---|---|
| AX-87529120 | 7.58E-14 | 11 | 83806589 | Zinc finger BED domain-containing protein |
| ctg7180001921007_7683_SCT | 1.90E-12 | 11 | 18233181 | N-acetylgalactosaminyl transferase 18-like |
| AX-96443306 | 1.10E-19 | 12 | 15637088 | Tf2-1<br>Fibroblast growth factor receptor-like 1 |
| AX-87148554 | 1.04E-15 | 12 | 79387657 | Neuron navigator 1-like isoform X(1-16) |
| ctg7180001848546_3400_SCT | 9.94E-09 | 13 | 4828240 | Carbonic anhydrase 6-like<br>Collagen alpha-1(IV) chain-like |
| ctg7180001916197_181_SGT | 4.40E-08 | 14 | 84342640 | Pumilio homolog 1-like<br>sodium/potassium-transporting ATPase subunit beta-1-interacting protein 1 |
| ctg7180001830170_7262_SCT | 5.43E-11 | 15 | 74833173 | Kinesin like protein KIF1B,KIF1C |
| ctg7180001828593_3363_SCT | 2.10E-10 | 15 | 97165095 | glutamine-rich protein 2 –like<br>sortilin-like |
| ctg7180001808968_16150_SCT | 4.95E-10 | 15 | 38346370 | Nuclear receptor coactivator 7-like isoform X2<br>ubiquitin-protein ligase rnf146-like |
| ctg7180001815063_2825_SAG | 6.80E-11 | 16 | 70878146 | beta-1,3-galactosyltransferase 1-like |
| ctg7180001583653_971_SGT | 1.27E-08 | 16 | 31088398 | nucleotide-binding protein G(o) subunit alpha |
| ctg7180001913273_34722_SAG | 1.11E-12 | 17 | 47700898 | Forkhead box protein M1-like isoform X(1,2,3) |
| AX-87053692 | 5.71E-15 | 18 | 7057036 | DNA-directed RNA polymerase I subunit RPA2<br>Fidgetin-like protein 1<br>DNA-binding protein Ikaros-like Isoform X(1,2,3) |
| AX-87831528 | 1.86E-08 | 18 | 60054788 | Proline, glutamic acid and leucine-ricj protein 1-like isoform X(1,2,3)<br>Collagen alpha-6(IV) chain-like isoform X(1,2,3,4,5 |
| AX-96135963 | 2.68E-14 | 19 | 15646119 | Metalloproteinase-16 isoform X (1,2,3,4) |
| ctg7180001853683_8586_SCT | 7.96E-09 | 19 | 71734301 | Tf2-1 poliprotein |
| ctg7180001224033_5203_SCT | 9.19E-11 | 20 | 6388643 | 1-phosphatidylinositol 4,5-bisphosphate phosphodiesterase<br>eta-1-like |
| ctg7180001206239_480_SCT | 3.95E-09 | 20 | 78138041 | Amine-associated receptor 13c-like |
| AX-87774242 | 8.83E-14 | 21 | 51570271 | RCC2 homolog zinc finger |
| AX-87889001 | 9.67E-10 | 21 | 5030928 | Beta-lactamase-like 1 isoform X 1,2 |
| AX-96182427 | 7.93E-16 | 22 | 22403762 | Histone-lysine N-methyltransferase, H3 |
| AX-96304397 | 1.39E-09 | 23 | 31997187 | Transposase-like |
| AX-215429861 | 2.00E-09 | 23 | 8888152 | RIMS-binding protein 2-like |
| AX-87566402 | 2.74E-09 | 24 | 22770805 | Transposon Tf2-1 |
| ctg7180001883626_2847_SAC | 4.18E-08 | 25 | 23683473 | SOX-6-like |
| ctg7180001909158_5625_SAG | 1.31E-12 | 26 | 26242989 | Transposase-like |
| ctg7180001910456_2266_SAG | 7.70E-09 | 27 | 17472982 | vacuolar protein sorting-associated protein 26A |

| | | | | |
|---|---|---|---|---|
| AX-86903102 | 2.97E-09 | 28 | 27513051 | RNA-directed DNA polymerase homolog |
| AX-215406230 | 6.92E-08 | 29 | 40809955 | cytoplasmic protein NCK1-like |

**LD candidate genes**

| c | P-VALUE | CHR | POS | Protein |
|---|---|---|---|---|
| ctg7180001560779_174_SCT | 3.70E-12 | 1 | 143099916 | zinc finger BED domain-containing protein 5-like |
| ctg7180001664070_2019_SGT | 3.19E-08 | 1 | 802443 | attractin isoform X2 |
| AX-215441061 | 5.45E-12 | 2 | 54974339 | zinc finger MYM-type protein 1-like<br>52 kDa repressor of the inhibitor of the protein kinase-like |
| ctg7180001438242_810_SGT | 1.62E-08 | 2 | 22171557 | neural-cadherin-like |
| ctg7180001865965_4181_SAG | 9.75E-24 | 3 | 78352788 | transposon Tf2-1 polyprotein |
| ctg7180001838479_5521_SCT | 6.248e-22 | 3 | 72632031 | Coronin-1A<br>Claudin-4<br>serine/threonine-protein phosphatase alpha-2 isoform-like |
| ctg7180001816393_1520_SCG | 8.228e-22 | 3 | 86298251 | E3 ubiquitin-protein ligase rnf213-alpha |
| AX-87074389 | 2.511e-18 | 12 | 15608796 | alpha-2-macroglobulin-like<br>(EXON) |
| ctg7180001460681_1054_SAG | | 3 | 78223769 | Centrosomal protein of 112 kDa (Exon)** varios caen acá |
| AX-96174672 | 5.98E-11 | 4 | 54508208 | protein phosphatase 1E<br>rab5 GDP/GTP exchange factor isoform X3 |
| AX-87380935 | 5.81E-12 | 5 | 48998361 | zinc finger protein 236-like isoform X2 |
| AX-215422843 | 9.33E-10 | 5 | 6471903 | protein LDOC1L-like<br>protein FAM127-like |
| AX-87285911 | 3.21E-09 | 6 | 38184520 | kelch-like protein 11<br>synaptonemal complex protein SC65-like |
| AX-86933745 | 2.23E-08 | 6 | 77405961 | transposase-like<br>alpha-(1,6)-fucosyltransferase-like isoform X1 |
| ctg7180001473407_12699_SGT | 3.09E-08 | 7 | 41256458 | endoplasmin-like<br>transcription termination factor 1-like isoform X2 |
| ctg7180001930535_7810_SAG | 1.72E-12 | 9 | 35745395 | transposon Tf2-1 polyprotein |
| ctg7180001915213_4447_SAT | 1.65E-11 | 9 | 109502144 | olfactory receptor 8I2-like<br>spartin isoform X3 [Salmo salar] |
| AX-96495871 | 2.87E-15 | 10 | 66025311 | moesin-like<br>alpha-ketoglutarate-dependent dioxygenase alkB homolog 3 |
| AX-87046611 | 3.99E-13 | 10 | 2698859 | adhesion G protein-coupled receptor L2 isoform X17<br>latrophilin-3 isoform X14 |
| AX-87323716 | 1.13E-13 | 11 | 58923059 | microtubule-associated serine/threonine-protein kinase 2-like isoform X7 |
| ctg7180001921007_7683_SCT | 4.32E-13 | 11 | 18233181 | polypeptide N-acetylgalactosaminyltransferase 18 isoform X1 |

| | | | | |
|---|---|---|---|---|
| | | | | zinc transporter SLC39A7 isoform X1 |
| AX-96443306 | 8.79E-20 | 12 | 15637088 | transposon Tf2-1 polyprotein<br>fibroblast growth factor receptor-like 1 isoform X1 |
| AX-87148554 | 3.47E-12 | 12 | 79387657 | neuron navigator 1-like isoform X13 |
| AX-87986193 | 2.33E-12 | 13 | 50890593 | transposase-like<br>fibroblast growth factor receptor-like 1 isoform X1 |
| AX-87545045 | 1.36E-09 | 14 | 25234547 | probable G-protein coupled receptor 141<br>transposase-like |
| AX-87361598 | 8.89E-11 | 15 | 72752181 | PHD finger protein 20-like |
| ctg7180001917967_1924_SAG | 1.04E-10 | 15 | 45943546 | zinc finger protein 2 homolog |
| ctg7180001315840_691_SGT | 5.92E-09 | 15 | 10204303 | E3 ubiquitin/ISG15 ligase TRIM25-like<br>tripartite motif-containing protein 47-like |
| AX-88146011 | 2.68E-08 | 16 | 81966618 | interleukin-1 receptor accessory protein-like 1-B |
| ctg7180001815508_106_SGT | 6.93E-11 | 17 | 30331957 | actin cytoskeleton-regulatory complex protein pan1-like |
| AX-87053692 | 1.36E-13 | 18 | 7057036 | fidgetin-like protein 1<br>DNA-directed RNA polymerase I subunit RPA2 |
| ctg7180001925448_27391_SGT | 4.67E-09 | 18 | 47165336 | transposase-like<br>neurexin-2-like isoform X1 |
| AX-215412266 | 3.75E-13 | 19 | 20059526 | general transcription factor II-I repeat domain-containing protein 2A-like<br>zinc finger BED domain-containing protein 5-like<br>neurexin-2-like isoform X12 |
| AX-96454132 | 6.38E-10 | 19 | 52344365 | protein RCC2 homolog |
| ctg7180001224033_5203_SCT | 1.18E-09 | 20 | 6388643 | transposon Tf2-1 polyprotein<br>protein RCC2 homolog |
| ctg7180001929365_7212_SAG | 7.68E-09 | 20 | 13375107 | mevalonate kinase isoform X1<br>cob(I)yrinic acid a,c-diamide adenosyltransferase, mitochondrial |
| AX-87889001 | 1.82E-12 | 21 | 5030928 | transposase-like |
| AX-87033709 | 1.90E-11 | 21 | 52176431 | protein RCC2 homolog<br>olfactory receptor 11A1-like |
| AX-87294473 | 2.36E-15 | 22 | 22996672 | zinc finger BED domain-containing protein 4-like |
| AX-87100741 | 4.08E-08 | 23 | 2081829 | RIMS-binding protein 2-like |
| AX-87566402 | 2.15E-09 | 24 | 22770805 | alpha-ketoglutarate-dependent dioxygenase FTO-like |
| AX-215432822 | 3.57E-10 | 26 | 29728927 | transposase-like<br>ZK1236.4 |
| ctg7180001910456_2266_SAG | 3.07E-09 | 27 | 17472982 | vacuolar protein sorting-associated protein 26A |
| AX-86903102 | 1.34E-08 | 28 | 27513051 | gamma-crystallin M2-like<br>delta-1-pyrroline-5-carboxylate synthase-like isoform X1 |
| ctg7180001888874_8563_SAC | 3.88E-08 | 28 | 2828690 | protein LDOC1L-like |

| | | | | |
|---|---|---|---|---|
| AX-87630043 | 3.39E-09 | 29 | 702591 | phenylalanine--tRNA ligase, mitochondrial-like |
| AX-86948954 | 7.64E-09 | 29 | 8888417 | SCAN domain-containing protein 3-like |

**LOG LD candidate genes**

| SNP | P-VALUE | CHR | POS | Protein |
|---|---|---|---|---|
| ctg7180001560779_174_SCT | 1.21E-11 | 1 | 143099916 | phenylalanine--tRNA ligase, mitochondrial-like |
| ctg7180001473327_1142_SAG | 9.71E-10 | 1 | 24091173 | protein LDOC1L-like<br>partitioning defective 3 homolog isoform X1 |
| ctg7180001194015_518_SGT | 4.84E-11 | 2 | 46130886 | probable G-protein coupled receptor 75 |
| ctg7180001816393_1520_SCG | 2.45E-31 | 3 | 86298251 | Timp2 Metalloproteinase inhibitor 2<br>GDP-L-fucose synthetase<br>transposon Tf2-1 polyprotein<br>fibroblast growth factor receptor-like 1 isoform X1 |
| ctg7180001919088_9536_SGT | 9.17E-10 | 3 | 8680933 | leucine-rich repeat-containing G-protein coupled receptor 4-like<br>transcription factor PU.1-like<br>myosin-binding protein C, cardiac-type-like isoform X6 |
| AX-215436146 | 1.75E-09 | 4 | 63749525 | zinc finger BED domain-containing protein 5-like |
| AX-87380935 | 3.87E-15 | 5 | 48998361 | transposon Tf2-1 polyprotein<br>RNA-directed DNA polymerase homolog<br>fibroblast growth factor receptor-like 1 isoform X1 |
| AX-215422843 | 4.32E-10 | 5 | 6471903 | transposon Tf2-1 polyprotein |
| ctg7180001877771_4032_SCT | 2.13E-10 | 6 | 21804607 | zinc finger BED domain-containing protein 4-like<br>fructose-bisphosphate aldolase A isoform X1 |
| AX-86921813 | 2.03E-08 | 6 | 55564139 | protein RCC2 homolog |
| ctg7180001802903_7539_SCT | 8.94E-10 | 7 | 45014082 | protein RCC2 homolog |
| ctg7180001820356_553_SGT | 7.63E-16 | 9 | 119887281 | clumping factor B-like, partial<br>potassium voltage-gated channel subfamily E member 4<br>E3 ubiquitin-protein ligase TRIML1 |
| ctg7180001378306_2288_SCT | 3.72E-14 | 9 | 39106556 | transposon Tf2-1 polyprotein<br>RNA-directed DNA polymerase homolog |
| ctg7180001574613_2026_SCT | 4.49E-13 | 9 | 128364943 | RNA-directed DNA polymerase homolog<br>moesin-like |
| AX-96495871 | 3.83E-13 | 10 | 66025311 | neurexin-2-like isoform X(1-13) |
| AX-87046611 | 5.61E-13 | 10 | 2698859 | cyclin-J<br>coiled-coil and C2 domain-containing protein 2A-like isoform X2 |
| AX-87284644 | 3.29E-16 | 11 | 84156626 | transposon Tf2-1 polyprotein<br>RNA-directed DNA polymerase homolog |

| | | | | |
|---|---|---|---|---|
| | | | | moesin-like |
| ctg7180001898739_2235_SAG | 4.90E-12 | 11 | 41716645 | oocyte zinc finger protein XlCOF8.4-like |
| ctg7180001873236_7461_SCT | 1.57E-11 | 11 | 16954030 | transposon Tf2-1 polyprotein<br>RNA-directed DNA polymerase homolog, partial |
| AX-96443306 | 5.73E-19 | 12 | 15637088 | RNA-directed DNA polymerase homolog |
| AX-86949583 | 2.80E-14 | 12 | 31185795 | probable G-protein coupled receptor 141 |
| AX-87265895 | 1.75E-12 | 12 | 78162841 | cohesin subunit SA-2-like<br>THO complex subunit 2-like |
| ctg7180001926541_2110_SCT | 1.79E-11 | 13 | 78971543 | protein phosphatase 1H<br>pleckstrin homology domain-containing family H member 1-like |
| AX-87545045 | 8.97E-11 | 14 | 25234547 | 28S ribosomal protein S28, mitochondrial<br>zinc finger BED domain-containing protein 5-like<br>calpain-1 catalytic subunit-like |
| ctg7180001823698_2837_SGT | 6.56E-12 | 15 | 74529588 | transposon Tf2-1 polyprotein<br>RNA-directed DNA polymerase homolog |
| ctg7180001848943_198_SAC | 3.92E-10 | 15 | 42610168 | sterile alpha motif domain-containing protein 9-like<br>transposon Tf2-1 polyprotein<br>moesin-like |
| ctg7180001501848_683_SAG | 2.42E-08 | 16 | 67652319 | homeodomain-interacting protein kinase 3-like isoform X4 |
| ctg7180001819429_4726_SAC | 9.23E-11 | 17 | 48666693 | SH2 domain-containing adapter protein F-like isoform X9<br>G2/mitotic-specific cyclin-B1-like isoform X2<br>ubiquitin-conjugating enzyme E2 Q2-like isoform X2 |
| AX-87009757 | 4.45E-14 | 18 | 7658606 | alpha-ketoglutarate-dependent dioxygenase alkB homolog 3<br>moesin-like |
| ctg7180001925448_27391_SGT | 2.65E-10 | 18 | 47165336 | adhesion G protein-coupled receptor L2 isoform X17<br>latrophilin-3-like, partial |
| ctg7180001233458_2510_SAG | 1.86E-13 | 19 | 23335967 | agglutinin-like protein 10, partial<br>protein RCC2 homolog |
| ctg7180001866830_1884_SCT | 1.51E-11 | 19 | 68273335 | zinc finger BED domain-containing protein 5-like<br>protein RCC2 homolog |
| ctg7180001192662_2876_SCT | 7.23E-10 | 20 | 67585936 | pleckstrin homology domain-containing family H member 1-like<br>transposon Tf2-1 polyprotein |
| AX-87889001 | 3.64E-15 | 21 | 5030928 | semaphorin-3C-like<br>platelet glycoprotein 4-like |
| AX-87774242 | 1.24E-13 | 21 | 51570271 | protein THEMIS-like<br>receptor-type tyrosine-protein phosphatase kappa-like isoform X2 |
| AX-87294473 | 2.97E-13 | 22 | 22996672 | inter-alpha-trypsin inhibitor heavy chain H3-like isoform X2<br>general transcription factor IIF subunit 1-like |
| AX-87100741 | 6.59E-10 | 23 | 2081829 | zinc finger protein 236-like isoform X2 |

| | | | | |
|---|---|---|---|---|
| | | | | protein RCC2 homolog |
| AX-96146224 | 1.71E-09 | 24 | 2125976 | protein LDOC1L-like<br>protein FAM127-like |
| AX-87864636 | 2.32E-08 | 25 | 23278154 | SH3 and PX domain-containing protein 2B isoform X4<br>tripartite motif-containing protein 16-like |
| AX-86930189 | 1.06E-10 | 26 | 25456427 | ubiquitin carboxyl-terminal hydrolase 36 isoform X3<br>metalloproteinase inhibitor 2-like |
| ctg7180001847078_1986_SGT | 1.21E-10 | 26 | 13287063 | exostosin-like 2<br>vascular cell adhesion protein 1-like isoform X2 |
| ctg7180001796728_5923_SCT | 2.98E-09 | 27 | 2705004 | zinc finger protein Aiolos-like<br>polycomb complex protein BMI-1-like<br>insulin-like growth factor binding protein 4 precursor<br>protein AF-17-like isoform X1 |
| ctg7180001794047_170_SGT | 1.70E-09 | 28 | 24356229 | zinc finger BED domain-containing protein 5-like<br>ribosome-binding protein 1-like |
| AX-87630043 | 5.99E-10 | 29 | 702591 | latent-transforming growth factor beta-binding protein 2-like<br>protein lin-52 homolog |
| AX-86948954 | 8.66E-10 | 29 | 8888417 | phenylalanine--tRNA ligase, mitochondrial-like |
